# The discovery of antimicrobial peptides from the gut microbiome of cockroach *Blattella germanica* using deep learning pipeline

**DOI:** 10.1101/2024.02.12.580024

**Authors:** Sizhe Chen, Huitang Qi, Xingzhuo Zhu, Tianxiang Liu, Yingda Teng, Qiuyu Gong, Cangzhi Jia, Tian Liu, S Chen, H Qi, X Zhu, T Liu, Y Teng, Q Gong, C Jia, T Liu

**Author notes:** Co-First Authors, these authors contributed equally to this study (S Chen, H Qi, X Zhu). **Corresponding Author: Tian Liu:**. E-mail address, **Cangzhi Jia:**. E-mail address, **Qiuyu Gong:**.

## Abstract

Antimicrobial peptides (AMPs) are candidates for use against antibiotic-resistant microorganisms. However, due to high cytotoxicity and poor performance in biological contexts, most AMPs are unable to satisfy the requirements. The gut microbiomes of pathogenic species like *Blattella germanica* represent unexploited reservoirs of naturally evolved biocompatible AMPs. Here we developed a lightweight AI pipeline called AMPidentifer with two nine-layers Dense-Net blocks and one embedded new self-attention module to enable the discovery of biocompatible AMPs from microbiome. The core structure of AMPidentifer is simple, not requiring complexing code basis. On the independent test dataset, it showed robust performance and avoided high false-positive results. From the gut microbiome of *B. germanica*, new AMP candidates with potential low toxicities and antimicrobial activities were identified by AMPidentifer. The selected two AMPs demonstrated good antimicrobial effects *in vitro* and *in vivo*. The Cys residue was demonstrated to perform different action mechanisms in the antimicrobial activity of two AMPs, providing insights for future rational design. New efficient AMPs with low cytotoxicity identified by the new high-throughput AI pipeline from the gut microbiome of *B. germanica* successfully showed an important interdisciplinary strategy for discovering bio-safe AMPs from nature.

The Graphic Abstract

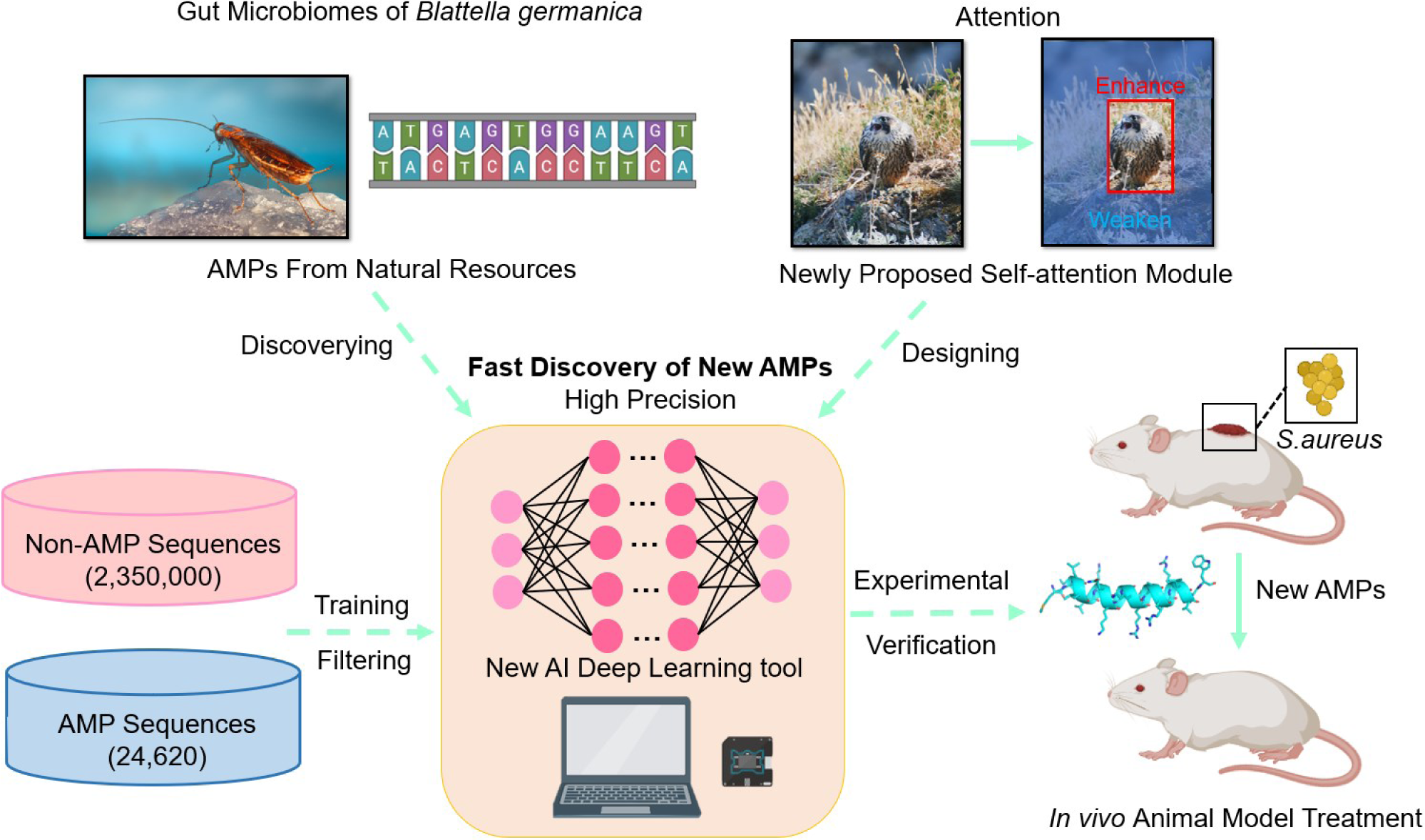

## Introduction

The wide applications of antibiotics have caused the quick spread of drug-resistant pathogens all over the world ^1–3^. Additionally, the ability to discover new antibiotics has been severely restricted, with only a small number of them becoming available for purchase recently. ^4^ Therefore, it is critical to create new strategies to combat microbial infections. Antimicrobial peptides (AMPs), which play a key role in regulating host immune responses and preserving the stability of microbial populations in hosts ^5–6^, are one interesting contender. Between 10 and 50 amino acid residues long, AMPs have an amphipathic and cationic property ^7^. Most cationic AMPs interact with negatively charged components of the microbial membrane, leading to membrane permeabilization and cell rupture ^8–9^. Only a small number of resistance mechanisms against AMPs can arise because the majority of AMPs typically kill microbes through non-specific physiochemical mechanisms ^10^. Additionally, AMPs often have substantially lower levels of environmental persistence ^11^. However, AMPs still have several problems with the application, e.g., low efficacy, degradations, or high cell toxicity. Only a small number of AMPs, including Omiganan, Surotomycin, and p2TA, have completed Phase III clinical trials ^12–13^. This highlights the limitations of traditional AMPs ^14–15^. Due to our inadequate understanding of structure-function correlations, it is difficult to use chemical alterations to increase effectiveness and reduce off-target conditions ^16^. A caveat is that such modifications may also increase risks of inappropriate inflammatory responses *in vivo* because many of the modified AMPs are derived from endogenous host defense peptides, which play crucial roles in modulating inflammatory responses in addition to their antimicrobial activities ^17^.

For a long time, various works have shown that it is very hard to achieve a counterbalance between low off-target cytotoxicity, good bio-compatibility, and potent antimicrobial effects *in vivo* by introducing unnatural amino acids or manually designed molecular blocks ^9^. The biological applicability of AMPs is frequently beyond the consideration of many synthetic chemistry laboratories, as many AMPs were specifically designed with low MIC values, which unfortunately lead to insurmountable host toxicities and failures of medical transformation during the past 30 years ^17–18^. However, compared to the limitations of human-involved chemical designs, seeking the naturally “designed” AMPs developed by the long history of biological evolutions might be more efficient and reliable. As these molecules only comprise L-amino acids, they can be easily synthesized by solid phase synthesis with low costs and limited byproduct reactions ^9^. For multi-cellular species like *Homo sapiens*, this is particular evidenced by such AMPs, retaining antimicrobial activity and biocompatibility, continusously act as a crucial roles in host-defence mechanisms for generation to generation ^18^. The existence of these AMPs indicated their low host toxicities and good biocompatibility in organisms ^19–20^. Various multi-cellular species harbor intrinsic gut microbial communities at different growth phases, which are actively necessary to hosts ^21^. The AMPs from the symbiotic gut microbes may specifically target invading microbes while avoiding toxicities to hosts; Otherwise, the hosts’ metabolism may be disrupted due to the off-target toxicities. That means, in theory, these unknown AMPs may be evolutionary meaningful and specifically target invading species and may be “designed” to evade bacterial resistance ^22–23^. This viewpoint was supported by new shreds of evidence ^24–26^. The human gut microbiome tended to exclude external pathogenic species and maintain the stabilities of the gut, indicating hypothetical specificities of these AMPs towards endogenous and exogenous microbes ^27–28^. Though at least 11 of the AMPs that are under clinical trials are derived from humankind ^5^, the possibility of the unfavorable cross-resistance of human gut microbiome to endogenous host AMPs is more concerning ^29^. Therefore, we focus on the gut microbiome of *B. germanica*, one of the most widely distributed pathogenic insects carrying harmful microbes like *Shigella Castellani*, *Pseudomonas aeruginosa*, *Salmonella enteritidis*, *Escherichia coli*, etc ^30–31^. *B. germanica* harbors harmful species without occurring pathogenesis ^32–33^. The consistent control of a variety of dangerous microorganisms in living systems suggested the potential presence of potent AMPs with low toxicity to hosts. In addition, their gut microbiome, which can be a useful resource for discovering novel AMPs, has never been reported for medical identification.

Until now, the identification of AMPs was still empirically driven by wet experiments in most cases ^9^. The experimental approaches are low-efficient, cost-intensive, and time-consuming ^5, 34^. Due to the gigantic predicating spaces, such identifications are mainly dependent on computational strategies. In recent years, the identifications of AMPs took advantage of machine learning (ML) and Deep Learning (DL) ^35–37^. These tools, whose AMPs prediction accuracy ranged from 87.17% to 92.11%, included support vector machine (SVM) ^38^, k-nearest neighbor (KNN), ^35^ random forests (RFs) ^39^, eXtreme Gradient Boosting (XGBoost) ^40^, deep neural network (DNN) ^41^, recurrent neural network (RNN) with long short-term memory (LSTM) ^42^, etc. ^43–44^. The rational design requirements for AMPs have not yet been satisfied, and novel AMP sequences are mainly derived from high-throughput space-searching ^45^. Most existing prediction models performed poorly in identifying non-AMP sequences, with only a 20-40% overall precision rate, indicating high false-positive ratios in real experiments ^35–44, 46^. Recently, a unified pipeline, incorporating 5 different complexing frameworks, was proposed to identify AMP from human gut microbiome data, and reached reasonable overall precision and specificities ^46^. However, their frameworks were at the risk of time-consuming, computationally-exhausting, and not user-friendly. Another tool HydrAMP is the first model designed to generate antimicrobial sequences including unconstrained and analogue generation methods ^47^. However, it was trained specifically for the task of analogue generation for the known AMP. In addition, the molecular dynamic (MD) simulations embedded in HydrAMP were time-consuming, expensive, and inaccurate. These features make it unsuitable for high-throughput screening of novel AMPs. Therefore, new user-friendly and simple pipelines are required to accelerate AMPs discovery while preventing false-positive conditions.

Here we proposed a pipeline called “AMPidentifer”, which showed high performance in predicting AMPs/non-AMPs (**Figure 1**). The core structure of AMPidentifer is simple, not requiring a complex code basis. And an attention module is mathematically designed in it, which reassigns the weights of every element in concatenated matrixes, potentially improving the recognition of key information in the input. AMPidentifer showed high performance in predicting AMP sequences, as measured by area under the precision-recall curve (AUPRC) and Precision. The source code and final model files are deposited at https://github.com/ChenSizhe13893461199/Fast-AMPs-Discovery-Projects. 79 candidate AMPs with new sequences were mined from the gut microbiome of *B. germanica* using AMPidentifer. By AI scores and other criteria for ensuring biosafety and effectiveness, two peptides (AMP 1 and AMP 2) were chemically synthesized. Both two peptides showed potent antibacterial activity and low mammalian cytotoxicity. AMP 1 even showed potent antimicrobial activities and wound-healing effects *in vivo*. Finally, the antimicrobial influences of single Cys residue in the two peptides were investigated, providing insights for future rational design.

**Fig. 1.**
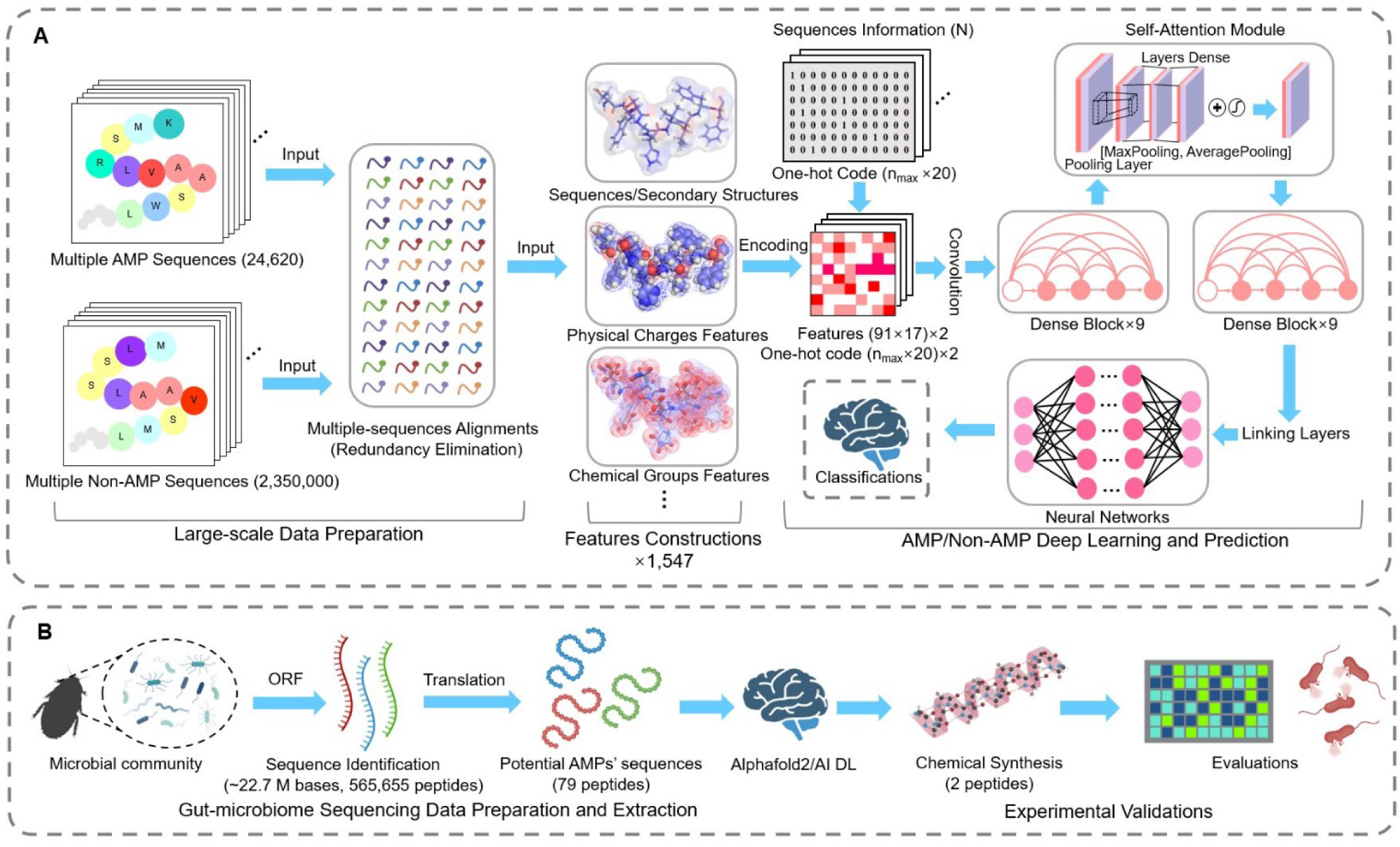
Schematic representation of AMPidentifer workflow. We mined gut microbiome data for potential AMPs, resulting in candidate AMPs for chemical synthesis and in vitro validations. **A**, Deep learning algorithms for identifying AMPs and non-AMPs, in which multiple features including physical, chemical, sequences, and structural information are utilized to construct input information for the model. **B**, AMPs mining strategies from gut microbiome data collected from pathogenic insect *B. germanica*.

## Result

### Designing AI deep learning strategy for efficient AMP Prediction

For input training data, the one-hot code and protein physicochemical descriptors were calculated to construct input representations for each peptide sequence. Sequence features of all selected AMP sequences were statistically analyzed (**Figure 2A-2B**), revealing pronounced amphipathic characteristics in their constitutions. Notably, among all amino acids, the positively-charged residues Lys (K) and Arg (R) possessed the highest distribution ratio while the negatively-charged residues Glu (E) and Asp (D) were relatively limited. This was hypothesized as the sequence prerequisite for most AMPs to function with killing effects because of the wide distribution of negative charges in most bacterial membranes. In addition, the hydrophobic residues like Leu (L), Val (V), and Ile (I) were also dominant, potentially contributing to the formation of amphipathic structures.

**Fig. 2.**
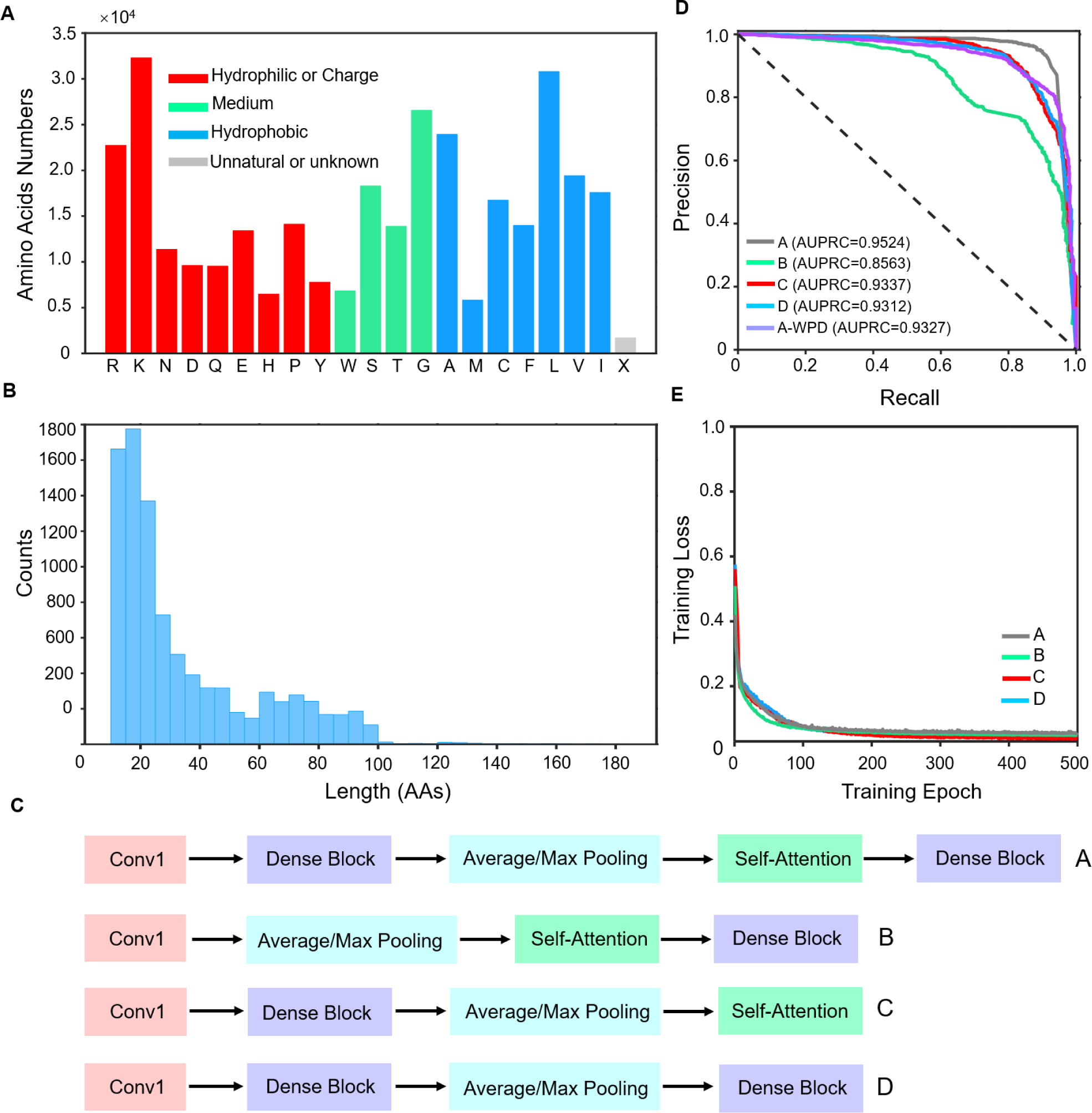
Establishing AMP prediction deep learning models. **A**, Amino acids distribution of AMPs collected from various databases for model training. X represents unknown or unconventional amino acids. **B**, the Length distribution of AMPs collected from various databases. **C**, Summary of four model strategies for testing and building. **(D)**, AUPRC evaluations for different combinations of strategies A, B, C, D, and A-without protein descriptors (A-WPD). **(E)**, Training Loss and circulation periods for different combinations strategies.

Here we designed a new DL pipeline to establish AMP identification tools, including two nine-layers densely connected convolutional neural network (Dense-Net) blocks and one embedded new self-attention module proposed in this work (**Figure 1**). Our basal model is an attention-based Dense-Net, where the original Dense-Net ^48^ is well renowned for low scales of fitting parameters and avoidance of gradient vanishing. The selection of model parameters and all hyperparameters are well illustrated in the **Materials and Methods**. For selecting the tool with the lowest false-positive ratio, our group optimized the performance of the model for potential AMPs discovery and primarily focused on the indicators of Precision and AUPRC. By iterative training and comparisons, we obtained the highest/average performance results of different model strategies (indicated as **A**, **B**, **C**, **D**) (**Figure 2C**). All evaluated model strategies converged rapidly within 500 epochs (**Figure 2C-2E**). For the highest performance results, the combinations of two nine-layer Dense-Net blocks, pooling layers, and newly designed self-attention module (**A**) showed the highest value of AUPRC (0.9524) and Precision (93.81%) (**Figure 2D** and **Table 1**). Interestingly, as shown in strategies **C** (AUPRC: 0.9337) and **D** (AUPRC: 0.9312), the introduction of the simple self-attention module showed a comparable performance to a nine-layers dense block. In addition, as shown in strategies **B** and **D**, only the matrixes preprocessed via a nine-layers dense block can be well recognized by the self-attention module and showed good performance (**Figure 2D** and **Table 1**). These results co-indicted the effectiveness of introducing self-attention modules following dense blocks. The average performance of different model strategies can also be found in the supplementary materials, and are consistent with this result (**Table S1**). We assumed that the self-attention module may improve the recognition of critical features and prevent information loss during training.

**Table 1.**
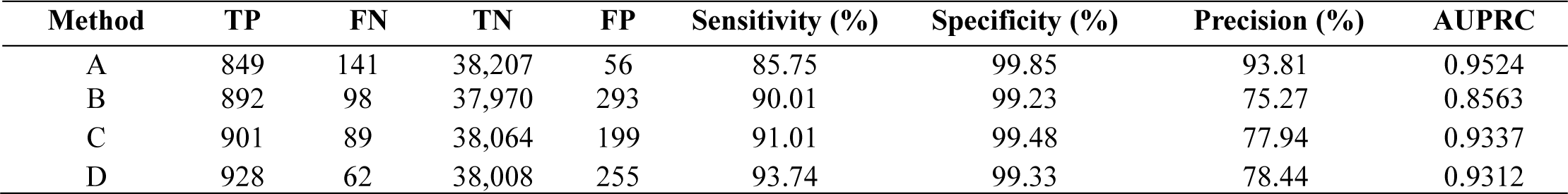
Comparison of the highest performance of different model designing strategies.

The representations of input data were also assessed. The combinations of one-hot code and protein physicochemical descriptors matrixes showed a higher AUPRC value (0.9524). It indicated the necessity of introducing physicochemical features, compared to the result of only using one-hot code input (AUPRC value of A-WPD: 0.9327) (**Figure 2D**). Such designs may increase the specificity of identification because the antimicrobial potency of AMPs was mainly based on physiochemical properties behind sequence compositions. It maintained low false-positive circumstances (Precision 93.81%), while retaining balancing Sensitivity at 85.75% and Specificity at 99.85%. Therefore, we selected structure A with the combined input of one-hot code and physiochemical descriptors as our final strategy for discovering new AMPs from microbiome.

For evaluating our pipeline among other frequently-used prediction algorithms, we trained these algorithms by the same training dataset of this work and obtained their highest/average performances by iterative training and comparisons (**Table 2** and **Table S2**). Ranked by the highest Precision and AUPRC indicators, our pipeline showed the highest performance among other tools (**Figure 2D** and **Table 2**). The average performance data of these tools can be found in the supplementary materials, and are also consistent with this tendency (**Table S2**). These results showed that our pipeline is a promising tool with a high degree of precision and low false-positive ratios.

**Table 2.**
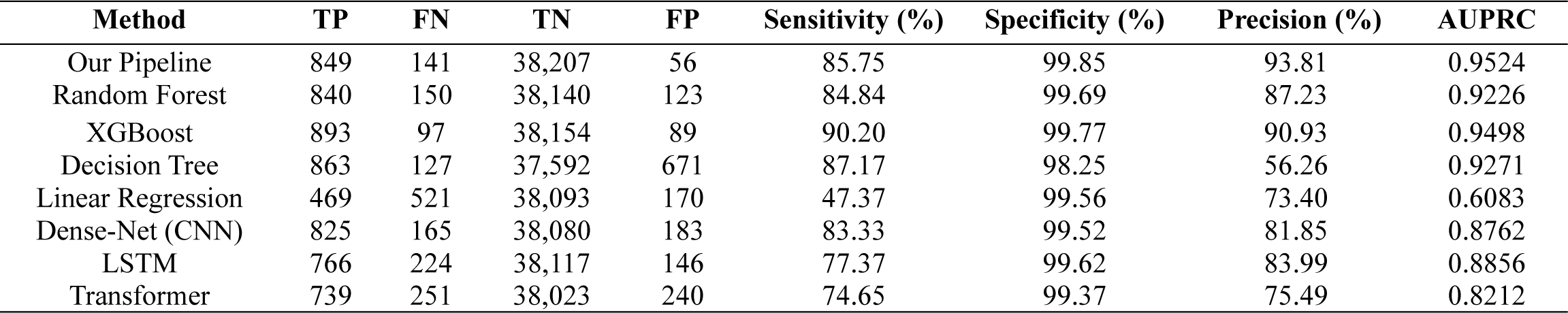
Comparison of the highest performance among frequently-used AMPs prediction algorithms^8–9, 35–46^.

### Mining potential AMPs from the gut microbiome of *B. germanica*

The gut microbiome of the pathogenic insect *B. germanica* possess an intricate community structure^49^. It is theoretically possible that the inner microbes could employ AMPs to compete for resources or dynamically stabilize the whole community structure. Based on available data of microbiome sequences of *B. germanica*, the sequential mining and subsequent filtering of potential AMPs were implemented to generate the putative AMP list. By filtering ∼22.7 M bases from 12 samples belonging to 6 growing stages of *B. germanica* (**Figure 1B** and **Figure 3A**), 565,655 putative peptides were extracted by open reading frame (ORF) identification, and 79 peptides were ultimately predicted as putative AMPs with potential possibility scores (PSP) larger than 50%. The single frequencies and overall occurring frequencies of these 79 peptides in different growth stages of *B. germanica* were shown in **Figure 3A**, suggesting their shared distribution features and potentially crucial functions.

**Fig. 3.**
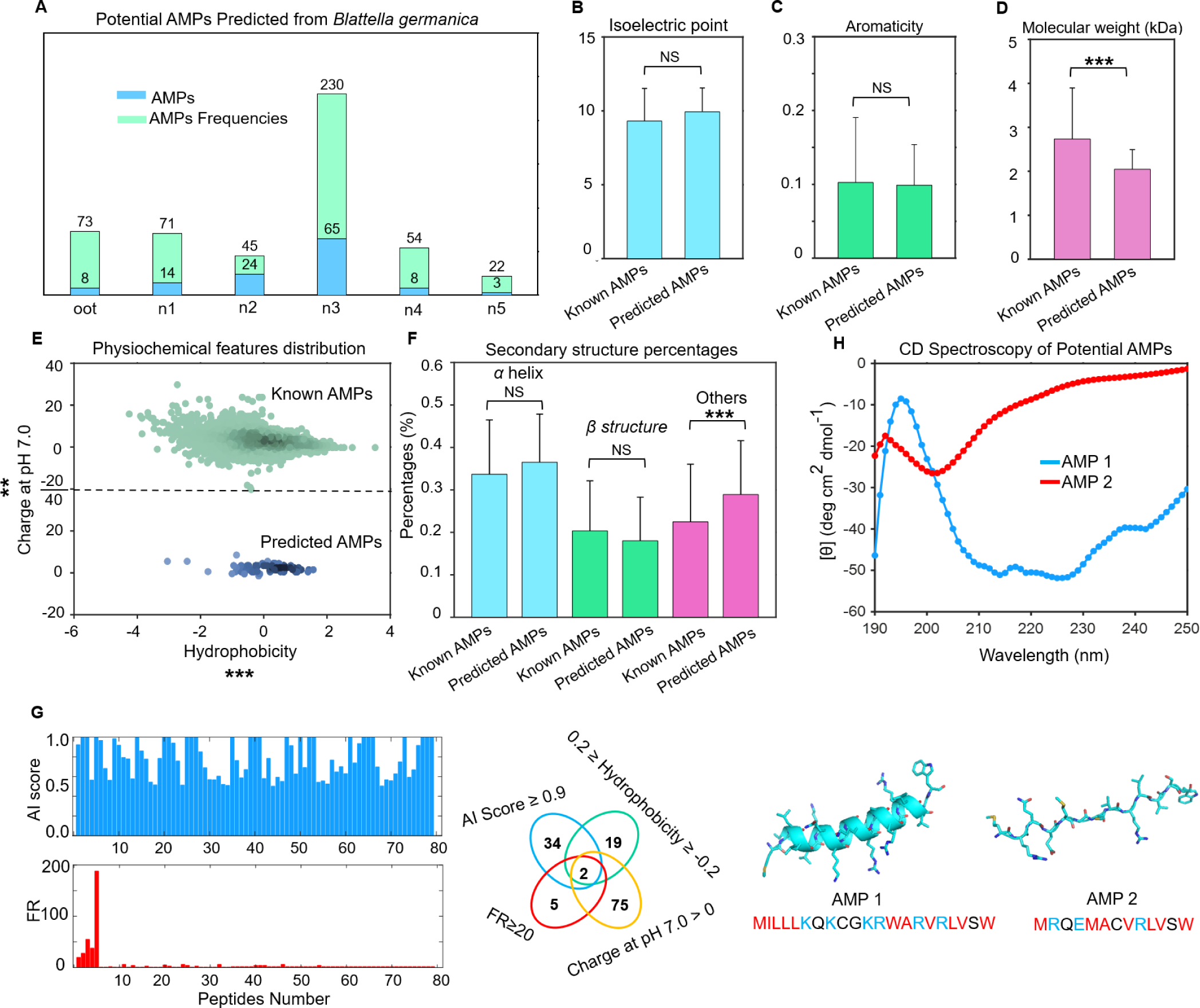
Mining potential AMPs from the gut microbiome of *B. germanica*. **A**, The statistics of potential AMP sequences predicted by the AI DL model from different growth stages (oot, n1, n2, n3, n4, and n5) of the gut microbiome of *B. germanica*. **B-F**, The comparison of physiochemical features of isoelectric points, aromaticity, molecular weight, hydrophobicity, charge at pH 7.0, and the secondary structure (*α* helix, *β* structure, and others) percentages between known AMPs and AI-predicted AMPs. **G**, the AI score and frequencies of 79 potential AMPs from *B. germanica*. Two sequences possessing AI scores> 0.9 and frequencies > 5, were further manually selected according to their sequential features for experimental validation. The structure of two peptide structures predicted by Alphafold2. **H**, The CD spectroscopy of secondary structures of two potential AMPs with concentrations of 0.2 mg/mL in deionized water.

The physiochemical features of isoelectric points, aromaticity, molecular weight, charge at pH 7.0, instability values, and the secondary structure percentages between these 79 peptides and the known AMPs were systematically analyzed (**Figure 3B-3F**). Notably, no significant differences in isoelectric points, aromaticity, and secondary structures were observed, indicating conserved characteristics among these features (**Figure 3B-3C** and **3F**). The distribution of molecular weights of AI-predicted AMPs showed smaller molecular weights compared to many of the known AMPs (**Figure 3D**). Many of the known AMPs possessed too many positive charges or even have negative charges at pH 7.0, suggesting their high toxicities or the lack of desired functions in biological environments. The charge distributions of AI-predicted AMPs (APA) at pH 7.0 mainly ranged between +1 and +9, with only 4 peptides possessing negative charge values (**Figure 3E**). These showed the balance between low toxicities and antimicrobial effects of cationic AMPs in living cockroach gut (pH≈7.0). The overall hydrophobicity of these APA convergently distributed around 0, suggesting conserved amphipathic patterns compared to many of the known AMPs. In summary, our AI pipeline showed potential capacities to identify sequences with the desired features.

9 peptides among the identified 79 peptides were found to have occurrence frequencies (OF) greater than 5 times, and 5 of them had OF between 20 and 188 times (**Figure 3G**). These 79 peptides were further assessed for AI scores, OF, charge values at pH 7.0, and hydrophobicity properties to ensure biosafety and efficacy. For the last round of experimental tests, 2 new peptides meeting the aforementioned criteria were chemically synthesized (**Figure 3G**). Notably, the first Met residues in these two possible AMPs were conserved because the methionine aminopeptidase lacks hydrolytic activity when the P1 position is occupied by big residues like Ile (I), Tyr (Y), Glu (E), and Arg (R).

The predicted structures of the two AMPs by Alphafold2 ^50^ showed amphipathic features along their chains, which are the prerequisites for most AMPs to show antimicrobial activities. The experimental results of CD spectroscopy supported the predicted structures of Alphafold2 (**Figure 3H**). While the absorbances at ∼200 nm in AMP 2 consistently demonstrated dominant random coils, the two negative peaks in CD spectroscopy of AMP 1 at 210 nm and 220 nm suggested the dominating helical structure. Both AMP 1 and AMP 2 were examined in the subsequent inhibition studies to determine their biological activities.

### Experiments verify the antimicrobial potencies of functional AMPs

The two peptides called AMP 1 and AMP 2 were chemically synthesized for verification and characterization. These two peptides displayed relatively higher proportions of Lys (K), Arg (R), and Trp (W) residues and fewer Cys (C) and Met (M), possessing positive charges of +6 and +1, respectively. The antibacterial activity of the two peptides against *S. aureus 2904*, *B. subtilis 23857*, *E. coli DH5α,* and *P. aeruginosa 2512* were tested at 200 μM concentration in liquid media. These two peptides were verified with antibacterial activity to at least one of these aforementioned four bacteria (**Figure 4A**).

**Fig. 4.**
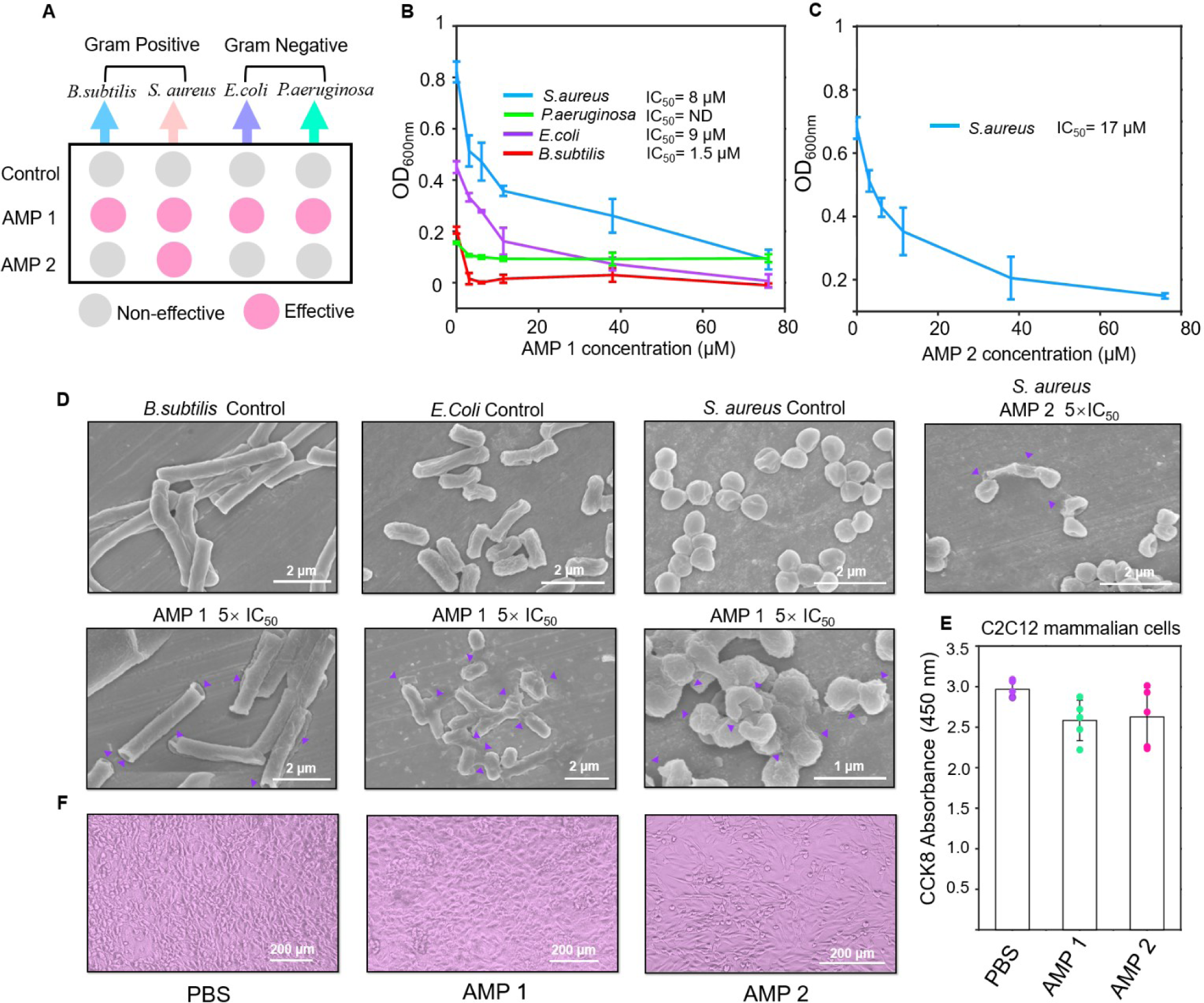
Experiments verify the strong potencies of functional AMPs. **A**, The screen of bacterial inhibition of four peptides against *S. aureus*, *B. subtilis*, *E. coli DH5α,* and *P. aeruginosa* at the concentration of 200 μM. **B**, The corresponding IC_50_ values and inhibition effects of AMP 1 and **C,** AMP 2 towards different bacteria at concentrations of 0 μM, 3.8 μM, 6 μM, 11 μM, 38 μM, and 76 μM cultivating for 7-8 h, respectively. **D,** The SEM examination of *Staphylococcus aureus*, *Bacillus subtilis*, and *Escherichia coli DH5α* cells treated with AMPs, showed cell content leakage and disruption of cell wall/membrane. All ruptured cells were indicated by a purple arrow. **E,** The C2C12 mammalian cells with 0.5% PBS solution or 100 μM AMPs were cultivated for 15 h at 37 °C. The cell activities and survival rates were indicated by the absorbance of CCK8 at 450 nm with 5 independent replicates (average survival rates=87±8.4% and 88.5±12.3%, respectively). **F,** the representative optical microscopy images of C2C12 cells.

At lower concentrations, AMP 1 and AMP 2 showed antimicrobial activities against *S. aureus 2904* with IC_50_ values of 8 μM and 16 μM, respectively (**Figure 4B-4C**, **Table 3**). Additionally, AMP 1 showed activity against *B. subtilis 23857* and *E. coli DH5α*, with IC_50_ values of approximately 1.5 μM and 9 μM, respectively. SEM was used to assess how AMPs 1 and AMP 2 affected these pathogenic bacteria because many AMPs kill bacteria through membrane rupture and cell lysis processes. The SEM results unmistakably showed that the cell membranes disintegrated and lysed at 5×IC_50_ concentrations of the relevant AMPs, confirming the deterioration of the cell membranes integrities (**Figure 4D**).

**Table 3.**
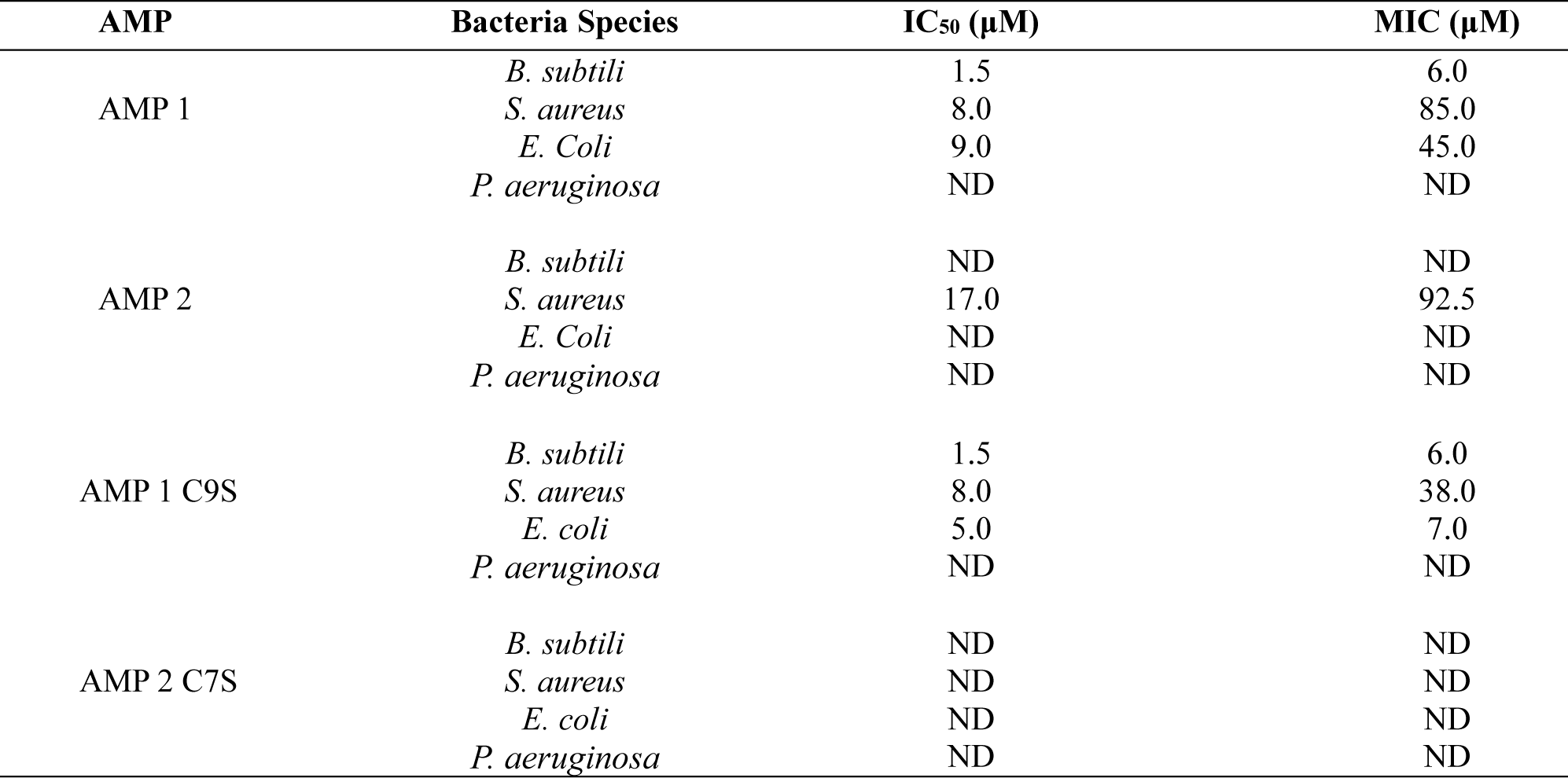
Antimicrobial activities of the identified AMPs and mutated sequences.

It has been concluded that the occurrence of resistance is unlikely because AMPs rarely involve specific molecular targets ^8^. With broad-spectrum efficiency, the AMP 1 tended to target gram-positive bacteria *S.aureus 2904*, *B.subtilis 23857*, as well as gram-negative species *E.coli DH5α*. While AMP 2 tended to specifically target gram-positive bacteria *S.aureus 2904*, avoiding the broad-spectrum killing of all microbes. The toxicities of these two cationic AMPs in mammalian cells were further investigated because off-target toxicity is a significant concern. At 100 μM high concentrations, both AMP 1 and AMP 2 exhibited low toxicities towards mammalian C2C12 cell activities (average survival rates=87±8.4% and 88±12.3%, respectively) (**Figure 4E-F**), indicating that their potential biocompatibility.

### The AMP 1 showed effective antimicrobial efficiencies *in vivo* animal model

Combining the aforementioned results, we selected the AMP 1 with better antimicrobial activities for conducting experiments *in vivo*. By using epithelial infection mouse models by pathogenic *S. aureus ATCC29213*, the infected mice group treated with the AMP 1 showed potent wound healing efficiencies, compared to the control group (**Figure 5A**). In addition, AMP 1 also revealed comparably curable effects with the commercialized antibiotic Vancomycin at the same dosage *in vivo*. Compared to the control group, the c.f.u. of the *S.aureus 2904* revealed decreasing tendencies after 12 days treatment of AMP 1 (**Figure 5B-5C**), showing comparably antimicrobial effects with Vancomycin. Tissue histology images of H&E and Masson showed that the tissues treated with AMP 1 or Vancomycin remained healthy morphology, while the control group showed severe tissue damage. To obtain more insights into the results, the bio-markers and inflammatory cytokines in tissues were revealed (**Figure S2**). Compared to the control group, the AMP 1 treatment group showed similar IL-1*β* levels at Day 12 without significant statistical differences (**Figure S2**), indicating that the AMP 1 may preferably function via direct antimicrobial effects, rather than the inflammation responses, to reveal therapeutic effects. In addition to IL-1*β*, the indicator levels of *α*-SMA, CD31, and FGF2 in AMP 1 are higher than the control group with significant statistical differences, indicating the cell growth and tissue recovery at infection sites. These experimental results suggested the potential clinic values of newly identified AMP 1 in antimicrobial applications.

**Fig. 5.**
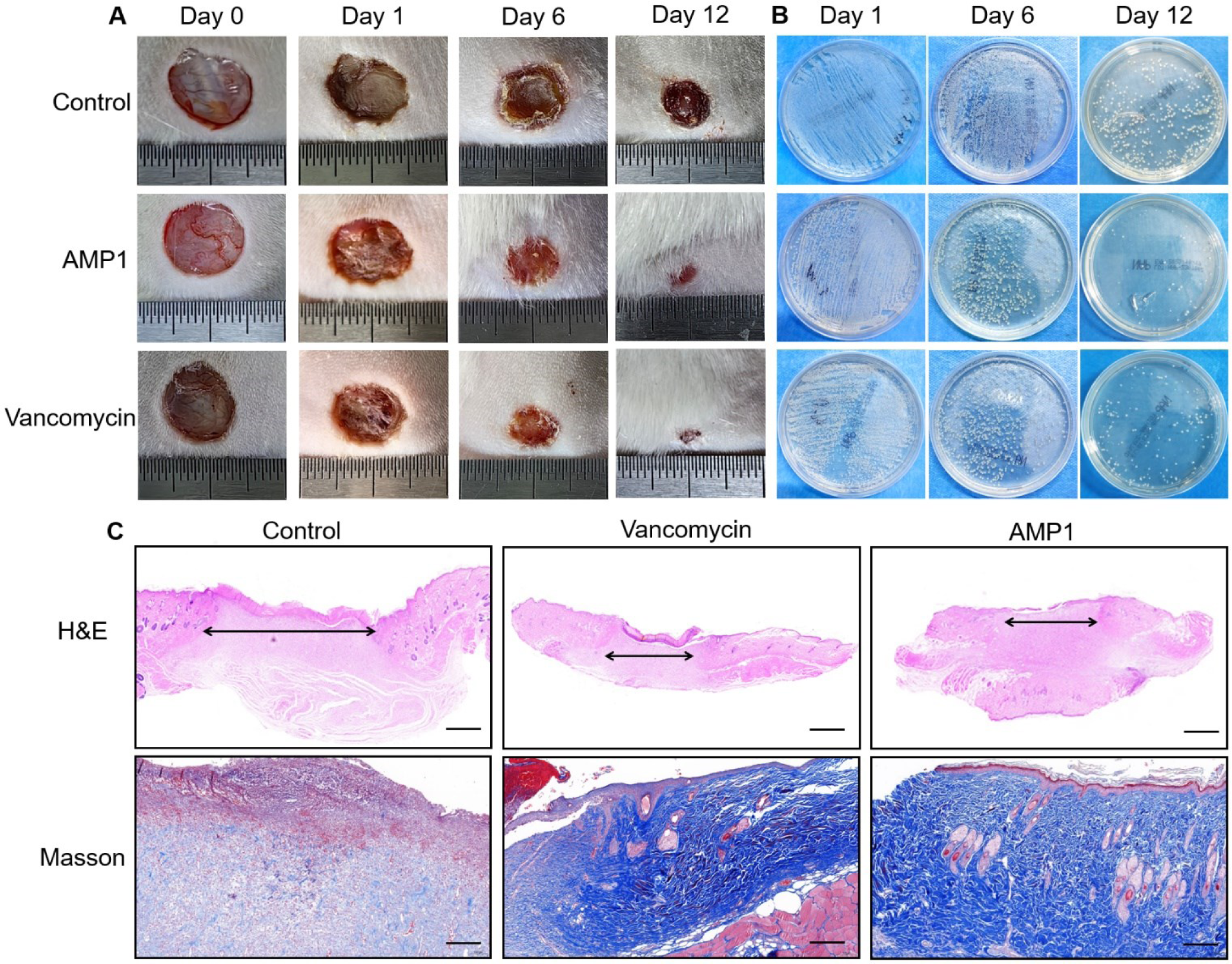
The antimicrobial and wound healing therapeutic effects of AMP 1 *in vivo*. **A,** the representative wound ealing results of the control PBS, Vancomycin, and AMP 1 to infected mice model at the same dosage *in vivo*. **B,** The xperimental verifications of infected *S.aureus* load of the control group, antibiotic Vancomycin treatment group, and MP 1 treatment group by streaking on LB agar medium plate. **C,** The representative histological images of infection ssues from the control group, Vancomycin treatment group, and AMP 1 treatment group generated by experimental &E and Masson dye methods

### The roles of single Cys residue in antimicrobial activities

Both AMP 1 and AMP 2 contained one Cys residue, which was potentially sensitive to oxidation and may form intermolecular disulfide bonds as reported in previous studies ^51–54^. Here, we attempted to explore the potential roles by mutating the Cys residue to the Ser residue.

The antimicrobial activities of AMP 1 C9S against *S. aureus 2904* and *B. subtilis 23857* were similar to those of the wild type, with IC_50_ values of 8 μM and 1.5 μM, respectively. The activities of AMP 1 C9S against *E. coli* showed improvement compared to the wild type, with an IC_50_ value of 5 μM and a MIC value of 7 μM. However, AMP 2 C7S exhibited a contrasting result with the disappearance of its antimicrobial activity (**Figure 6A-6B**, **Table 3**). These results indicated that the Cys residue probably performed different roles in AMP 1 and AMP 2.

**Fig. 6.**
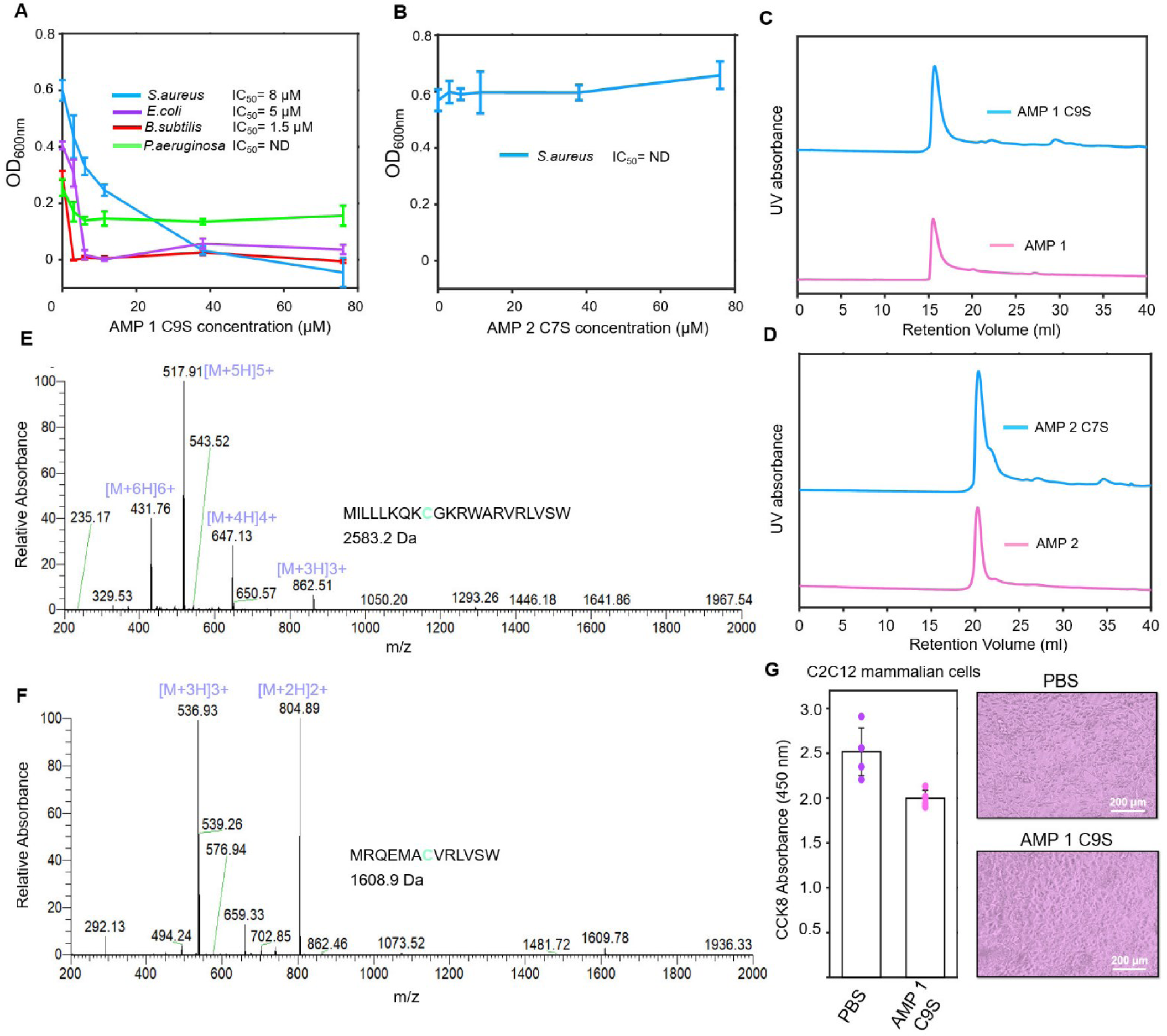
The influences of Cys residues in antimicrobial activities. **A,** AMP 1 C9S and **B,** AMP 2 C7S towards different pathogenic bacteria at concentrations of 0, 3.8, 6, 11, 38, and 76 μM at 37 °C cultivating for 7-8 h, respectively. The Mass spectrometry of **C,** AMP 1 and **D,** AMP 2 diluted in deionized water at the concentrations of 2 mg/mL. The SECresults of **E,** AMP 1and AMP 1 C9S, **F,** AMP 2and AMP 2 C7S at 25 °C in buffer containing 150 mM NaCl, 20 mM Tris, pH 7.0. **G,** The C2C12 mammalian cells with 0.5% PBS solution or 100 μM AMP 1 C9S were cultivated for 15 h at 37 °C. The cell activities and survival rates were indicated by the absorbance of CCK8 at 450 nm with 5 independent replicates (average survival rates=80±3.5%).

Then, we used MALDI-TOF mass spectrometry to detect potential dimers formed by disulfide bonds in AMP 1 or AMP 2 to better understand the molecular mechanism of the Cys residue. The results demonstrated that both peptides may perform antibacterial functions as monomers (**Figure 6E-6F**). To further validate these results, size-exclusion chromatography (SEC) was performed. The dominant presence of the monomer was demonstrated by the single peak appearing at the same retention volumes of 15 ml in both AMP 1 and AMP 1 C9S (**Figure 6C**). Similar results were also obtained from the SEC analysis of AMP 2/AMP 2C7S (**Figure 6D**). These results suggested that there was no existence of the intermolecular disulfide bonds in these two AMPs. Regarding AMP 1, the mutation of Cys residue to Ser residue not only increased its antibacterial activity, but also potentially improved the stability by decreasing the redox properties of the peptide, providing a strategy for the rational design of AMPs.

Additionally, as shown by CCK8 reagents, C2C12 mammalian cells treated with high concentrations of 100 μM AMP 1 C9S maintained high survival rates (average survival rates=80±3.5%) and cell activities (**Figure 6G**), which demonstrated its potential engineering values for future research.

## Discussion

In this study, we proposed a new computational pipeline called “AMPidentifer”, identifying two new AMPs from the gut-microbiome of cockroach *B. germanica*, which showed potent antibacterial activities and low mammalian toxicities. *In vivo*, the AMP 1 showed good wound-healing effects and potent antimicrobial activities similar to the antibiotic Vancomycin in lowering *S.aureus* load. Furthermore, we demonstrated that the antimicrobial activities of both peptides were not mediated by the formation of disulfide bonds between Cys residues.

The AMPidentifer showed high performance in precision and AUPRC. The lightweight features of the self-attention module made it possible to possess low-scale fitting parameters and avoid over-fitting conditions. Our pipeline overcame the challenges of high false-positive circumstances in predicting non-AMPs (99.85%) and had a high AUPRC value (0.9524). In addition, our pipeline identified AMPs with desired features (low toxicities, moderate hydrophobicity, and rational cationic charges), contributing to avoiding high toxicities under biological environments. ^12–15, 53^ Although the AMPs embrace a long history of medical research, the current AMPs deposited in libraries only account for a small fraction of all AMPs in nature. The discovery of AMPs from the gut microbiome of *B. germanica* has never been reported before this work. The applications of AMPs from the gut microbiome of *B. germanica*, rather than from the human gut microbiome ^34^, potentially decrease drug-resistant evolution risks of the human microbial community. The evolved resistance to human gut-derived AMPs may lead to undesirable resistance of endogenous microbial communities in the human body.

By new pipeline and gut microbiome data of *B. germanica*, we identified two new AMPs and one mutated AMP with antimicrobial activity comparable to that of previously reported AMPs with low cytotoxicity, such as c_AMP1043^46^, Olabogan^47^, Ratigan^47^, and CRRI3^55^. In infection mouse models by *S. aureus 2904*, the AMP 1 showed potent antimicrobial efficiencies and wound-healing effects *in vivo*. The sequences of AMP 1 and AMP 2 were unique with low shared sequential features with known AMP sequences, as well as single Cys residue contained in sequences. Previous studies have shown that the majority of AMPs are enriched with an even number of cysteines, which elicits the formation of multiple intramolecular disulfide bonds that are crucial to act as a chemical shield against foreign microorganisms, such as C4VG16KRKP ^51^, and vancomycin ^52^. Interestingly, disulfide bonds were not found in any of the two AMPs in this study. The AMP 2 C7S lost its activities. We unexpectedly found the increased antimicrobial activity of AMP 1 C9S, indicating the complexity of the action mechanism of Cys residue in the antimicrobial activity of peptides. Inspired by these results, we proposed that this residue has excellent potential for the future rational design of antimicrobial peptides.

### Conclusions

In summary, this study proposed the strategy of applying the new high-throughput AI tool to identify naturally evolved biocompatible AMPs from the unexploited gut microbiome of pathogenic *B. germanica*. The selected new peptides showed potent antimicrobial activities, microbial membrane destruction capabilities, and low mammalian cytotoxicity *in vitro* and *in vivo*. This work also highlights the significance of key residues in AMP sequences and provides further insights into the rational design of AMPs. The discovery of AMPs from the gut microbiome of *B. germanica* has not been reported before this work, and it represents a new interdisciplinary strategy for discovering naturally evolved bio-safe AMPs from nature.

## Experimental/Methods Section

### Data and code availability

The source codes are publicly deposited at https://github.com/ChenSizhe13893461199/Fast-AMPs-Discovery-Projects. This research contains publicly available AMP and non-AMP data. AMP data were collected from four public AMP datasets (ADAM ^55^, APD ^56^, CAMP ^57^, and LAMP ^58^), which contained most of the currently available AMPs from different resources (downloaded as of June 15^th^, 2022). The non-AMP dataset was downloaded from various resources deposited in UniProt by eliminating any entry that contains the following notes: antimicrobial, antibiotic, antiviral, antifungal, or effector (downloaded as of June 15^th^, 2022). The initial downloads contain 24,620 AMPs and 2,350,000 non-AMPs, which were downsized by multiple sequence alignment tool Cgwin with 90% and 50% cut-off threshold values, respetively. Lastly, only the 11 ≤peptides ≤50 amino acids (AAs) were considered since most AMPs are in this scope ^8^. Eventually, we got 8,792 AMPs and 79,675 non-AMPs for subsequent model construction. Based on the Needleman–Wunsch package in MATLAB R2021 a, the AMPs/non-AMPs were split as train/validation/test ratios of approximately 8:1:1 and 4.5:4.8:0.7, respectively. Such data division attempted to avoid the positive/negative data imbalance in model training. Therefore, the available sequences were split as the training dataset (6,998 AMPs and 36,406 non-AMPs) for model training, the validation dataset (804 AMPs and 5,006 non-AMPs) for hyperparameters fine-tuning, and the test dataset (990 AMPs and 38,263 non-AMPs) for final evaluations.

All codes were written by Python 3.9 and were implemented on servers with A40 (48 GB) GPUs. The highest/normal performance of each model strategy (A, B, C, and D) were determined via iterative training and comparisons. The highest/average performance data of each model strategy and the .h5 files of each model strategy can be found in the “Model” directory at the aforementioned GitHub link. Additionally, the .h5 files of the well-trained model A with the highest performance were deposited in a compressed file called “AMPfinder.rar. The Gut microbiome sequencing data of the omnivorous cockroach *Blatella Germanica*^49^ at different growth stages were selected to mine potential AMPs. The ORF prediction was implemented by a python script called “ORF_hunter.py” and it has been submitted to the GitHub link.

### Model structure design, hyperparameters and performance data deposition

For our pipeline, the optimal number of input matrixes (two 91×17 and two 20×50 matrixes, respectively), two nine-layers dense blocks, and two 1×7 and 1×11 filter kernel sizes were ultimately determined via model performance evaluations based upon the independent validation dataset. The initial learning rate was adjusted to 0.0015 with adaptive learning rate optimizer Adam and default parameters. The dropout rate and dropout density were all set as 0.2, with the batch sizes between 512 and 256. The weight decay was set as 1×10^-6^ and the growth rate of Dense-Net work was determined as 36. The training performance were repeatedly evaluated to rigorously verify each parameter listed here. Full details of hyperparameters can be found in the source codes at the aforementioned GitHub link. The trained model with the highest prediction performance was applied for further excavating AMPs from the gut microbiome data. Other standard algorithms including Random Forest, XGBoost, Decision Tree, Linear Regression, Dense-Net, LSTM, and Transformer were re-constructed for comparison, using the same training/validation dataset of this work to train and adjust hyperparameters, respectively. For aims of convenient accessibilities and fast utilizations, other previously proposed complexing frameworks were not considered here due to cumbersome structures, high computational-resources intensities, and time costs. The well-trained model files of each comparison model and their highest/average performance data can be found in the “Model” directory at the GitHub link.

### New Self-attention module design

Here we designed a new self-attention module, which was proposed for the first time and was demonstrated to be more effective than a nine-layers dense block in this work. The details of this module can be found in the supplementary materials (**Figure S1**), and the source codes with full parameters are available at the GitHub link.

### Protein descriptors

The AMPs/non-AMPs sequences were represented by one-hot matrixes and protein physicochemical descriptors (containing 1,547 different features). The one-hot matrix contained n_max_×20 elements (n_max_=50, representing the maximum amino acid numbers in all sequences) for describing amino acid compositions and sequential information.

The protein physicochemical descriptors contained 1,547 elements calculated by PyPro packages and were transformed to a 91×17 matrix (91×17=1,547), representing all currently available physicochemical features of peptide sequences including hydrophobicity, hydrophilicity, charges distributions, secondary structures, etc.

### Prediction pipeline standards

We defined several evaluation indicators to quantify the performance of different models. The Precision, Sensitivity (precision of AMPs prediction), and Specificity (precision of non-AMPs prediction), are defined as follows:

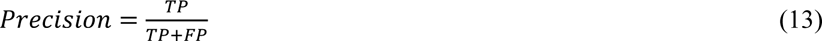

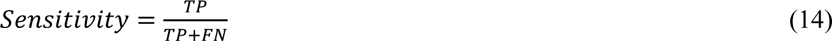

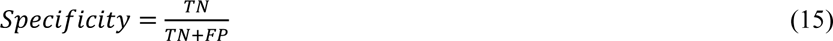

Where the TP, FP, TN, and FN indicated true positive number, false positive number, true negative number, and false negative number, respectively.

### Peptide synthesis

The peptides under investigation in this study were chemically synthesized using Gill Biotech’s solid-phase peptide synthesis. High-performance liquid chromatography was used to confirm that all peptides had purities that were higher than 95%. Mass spectrometry was used to confirm their molecular weights and sequence contents.

### Bacterial inhibition experiment and MIC determination

Four species of pathogenic bacteria including typical gram-positive bacteria *Bacillus subtilis* 23857 (*B. subtilis* 23857) and *Staphylococcus aureus* 2904 (*S. aureus* 2904), andtypical gram-negative bacteria *Escherichia coli DH5α* (*E. coli DH5α*) and *Pseudomonas aeruginosa* 2512 (*P. aeruginosa* 2512) were carefully streaked on Luriae–Bertani (LB) agar medium and incubated at 37 °C for 16 h. The individual colonies were further selected to cultivate in LB liquid medium and shaken at 120 rpm at 37 °C for 12 h. Then, the suspension was diluted to approximately 5×10^5^ colony-forming units (c.f.u.) per milliliter, for the final inhibition test and minimum inhibition concentration (MIC) calculations. We rapidly thawed the freeze-dried powder of AMPs and dissolved them in double-distilled water to a final concentration of 20 mg/ml. Following experimental procuderes of previously published works ^45–46^, we established three groups to verify the efficiencies of the four peptides: (1) blank control group, 200 μl of deionized water; (2) bacterial control group, 100 μl of deionized water and 100 μl of bacterial solution; and (3) peptides group (200 μM), 100 μl of peptides solutions (400 μM) and 100 μl of bacterial solution. For peptides showing apparent inhibition effects, our group further set 9 test groups cultivating for 7-8 h to detect AMP antibacterial activity: 0, 3.8, 6, 11, 38, 76, 85, 92.5, and 100 μM. All inhibition experiments were implemented on 96-well plates with each well containing a 200 μL volume sample (100 μL bacterial solution and 100 μL AMP solution). The OD_600_ values of each well were measured after culturing at 37 °C, 500 rpm with three independent replicates. MIC results were further verified by the spread plate method. The bacterial inhibition ratio was calculated via the formula:

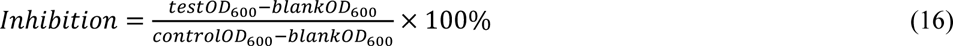

### Circular dichroism spectroscopy of potential AMPs

The secondary structures of all potential AMPs were identified using Circular dichroism (MOS-5000304040403, BioLogic Science Instruments, France) in deionized water at a fixed concentration of 0.2 mg/mL in a 1-cm quartz dish. The sample concentrations at 0.1, 0.2, 0.5, and 1 mg/mL were pre-tested and 0.2 mg/mL was selected to be the best concentration to avoid possible interferences to signals.

### SEM measurement

The membrane disruptions by AMPs were measured by scanning electronic microscopy (SEM). The *B.subtilis* 23857, *S. aureus* 2904, and *E.coli DH5α* were cultivated to the exponential phase at 37 °C. The bacterial suspensions (c.f.u. = 10^8^ ml^−1^) were treated with AMPs 1 and AMPs 2 at the concentrations of 5×IC_50_ at 37 °C for 24 h. Samples were washed with deionized water 3 times and further treated with 0.5 % glutaraldehyde for 3 h. The sample (10 µL) was added onto a copper wafer and then dehydrated with 75 % ethanol 5 times. The images were generated using SEM equipment (HITACHI S-4800, China).

### Cytotoxicity against mammalian cells

C2C12 murine myoblasts were purchased from the commercially available culture collection (RSBM). Cells were cultured and maintained in a 5 mL MEM medium containing 10% Fetal Bovine Serum (FBS) and 1% penicillin–streptomycin in an incubator at 37 °C, for 5 days under an atmosphere of 5% CO_2_ supply. Then, the cells were processed by pancreatic enzymes for 5 minutes and then centrifuged 1000×g for 5 min. The cell sediment was then resuspended in a 5 mL MEM medium and diluted to an ultimate density of ∼1×10^4^ cells/mL. The diluted cell suspensions were cultivated in 96-well plates (100 μL volume) for 2 days. Following experimental procuderes of previously published works ^45–46^, the tested groups were processed by adding 0.5% PBS, 100 μM AMP 1, or 100 μM AMP 2 for 15 h with five independent replicates. The cell survival rates were determined by adding 10 μL CCK8 in each well for 4 h and the absorbance of each well was determined at the wavelength of 450 nm. The average cell survival rates (%) were calculated based on the average absorbance differences at wavelength 450 nm between the PBS-control group and the AMPs-treated group.

### MALDI-TOF mass spectrometry

The peptides were diluted to 2 mg/mL by deionized water for further evaluation. The mass spectrometry detections of AMPs were consigned to the authorized agency at Dalian University of Technology, China. The mass spectrometry equipment was purchased from Thermo Fisher Scientific Inc. (Q Exactive Plus 20225220) with a resolution ratio of 280,000 FWHM and a nano-ESI source. The experiments were implemented in the m/z range from 200 to 2000. The following experimental conditions were chosen: sheath gas and auxiliary gas flow rates, 15 and 5 units; capillary temperature, 275 °C; spray voltage, 3.5 kV; in-source collision energy, 20.0 eV; and auxiliary gas heater temperature, 175 °C. The software Thermal Xcalibur Qual browser completed the final data analysis.

### The size-exclusion chromatography (SEC)

The SEC column applied in this work was Superdex 30 purchased from Cytiva Life Science. First, the SEC column was gently cleansed with deionized water for one hour at a flow rate of 0.6 mL/min. Then, at a flow rate of 0.6 mL/min, buffer solutions comprising 150 mM NaCl and 20 mM Tris, pH 7.0, were used to pre-equilibrate the SEC column. Finally, buffer-diluted AMPs were introduced into the column at a flow rate of 0.3 mL/min. UV absorbance and retention volumes were then used to determine the molecular sizes.

### *In Vivo* evaluation of antimicrobial efficiency

Healthy female SD rats (6-8 weeks, 180-220 g) were fully anesthetized and shaved on the back. A circular full-thickness wound of 10 mm in diameter on the back of the rat was removed by a hole puncher, and the wound was infected by S. Aureus (ATCC: 29213, 1 × 10^7^ CFU/mL, 100 μL) for 24 h. After 24 h of infection, all the mice were randomly divided into three groups (9 in each group): (1) PBS (control), (2) AMP1 (100 uM), (3) Vancomycin (100 uM). Then the rats were treated with 100 uL PBS, AMP 1, and vancomycin on days 1, 2, 3, 6, and 9. The wound healing processes were recorded by taking pictures on days 0, 1, 6, and 12. The rats were sacrificed (3 in each group) on days 1, 6, and 12, and the granulation tissue was divided into two parts. A tissue was immediately preserved for H&E, IHC, and Masson. Another tissue was placed in a sterile centrifuge tube with 2 mL PBS before being plated onto an LB agar plate to assess the bacterial colonies area. H&E, IHC, Masson, and Giemsa staining results were obtained from Servicebio, China.

### Ethics statement

All experiments involving animals were conducted according to the ethical policies and procedures approved by the Health Science Center of Xi’an Jiaotong University (New Permit Number: **XJTUAE2023-2129**).

## Supporting information

All supplementary materials and data

## Reference

1. E.Y. Klein, M. Milkowska-Shibata, K.K. Tseng, M. Sharland, S. Gandra, C. Pulcini, R. Laxminarayan Lancet. Infect. Dis. 2021, 21, 107–115. doi: 10.1016/S1473-3099(20)30332-7

2. H. Charles, M. Prochzaka, K. Thorley, A. Crewdson, D.R. Greig, C. Jenkins, A. Painset, H. Fifer, L. Browning, P. Cabrey, R. Smith, D. Richardson, L. Waters, K. Sinka, G. Godbole Lancet. Infect. Dis. 2022, 22(10), 1503–1510. doi: 10.1016/S1473-3099(22)00370-X

3. E.M. Darby, E. Trampari, P. Siasat, M.S. Gaya, I. Alav, M. A. Webber, J. M. A. Blair Nat. Rev. Microbiol. 2023, 21, 280–295. doi: 10.1038/s41579-022-00820-y

4. C. Årdal, M. Balasegaram, R. Laxminarayan, D. McAdams, K. Outterson, J. H. Rex, N. Sumpradit Nat. Rev. Microbiol. 2020, 18, 267–274. doi: 10.1038/s41579-019-0293-3

5. B.P. Lazzaro, M. Zaslof, J. Rolf Science. 2020, 368(6490), eaau5480. doi: 10.1126/science.aau5480

6. B. Kintses, O. Méhi, E. Ari, M. Számel, Á. Györkei, P. K. Jangir, I. Nagy, F. Pál, G. Fekete, R. Tengölics, Á. Nyerges, I. Likó, A. Bálint, T. Molnár, B. Bálint, B. M. Vásárhelyi, M. Bustamante, B. Papp, C. Pál Nat. Microbiol. 2019, 4, 447–458. doi: 10.1038/s41564-018-0313-5

7. W. Li, F. Separovic, N.M. O’Brien-Simpson, J. D. Wade Chem. Soc. Rev. 2021, 50, 4932–4973. doi: 10.1039/d0cs01026j

8. G.A.P. Aronica, L. M. Reid, N. Desai, J. Li, S. J. Fox, S. Yadahalli, J. W Essex, C.S. Verma J. Chem. Inf. Model. 2021, 61, 3172−3196. doi: 10.1002/psc.2947

9. D.T.T. Marcel, C. Jicong, L.F. Octavio, T.K. Lu, C. de la Fuente-Nunez ACS Nano 2021, 15, 2143−2164. doi: 10.1021/acsnano.0c09509

10. A. Peschel, H.G. Sahl Nat. Rev. Microbiol. 2006; 4(7): 529−536. doi: 10.1038/nrmicro1441

11. E. Gullberg, S. Cao, O.G. Berg, C. Ilbäck, L. Sandegren, D. Hughes, D. I Andersson. Selection of resistant bacteria at very low antibiotic concentrations. Plos. Pathogens. 2011, 7, e1002158. doi: 10.1371/journal.ppat.1002158

12. H.B. Koo, J. Seo. J. Pept. Sci. 2019, 111 (5), 24122. doi: 10.1002/pep2.24122

13. P.S. Gromski, A.B. Henson, J.M. Granda, L. Cronin Nat. Rev. Chem. 2019, 3, 119−128. doi: 10.1038/s41570-018-0066-y

14. J. Zhou, H. Zhang, M. S. Fareed, Y. He, Y. Lu, C. Yang, Z. Wang, J. Su, P. Wang, W. Yan, K Wang ACS NANO 2022, 14 (4), 758−766. doi: 10.1021/acsnano.1c11206

15. R. Mourtada, H.D. Herce, D.J. Yin, J. A. Moroco, T. E. Wales, J. R. Engen, L. D Walensky Nat. Biotechnol. 2019, 37 (10), 1186−1197. doi: 10.1038/s41587-019-0222-z

16. C. Loose, K. Jensen, I. Rigoutsos, R. Isidore, S. Gregory, Nature 2006, 443 (7113), 867−869. doi: 10.1038/nature05233

17. N. Mookherjee, M.A. Anderson, H.P. Haagsman, D.J. Davidson Nat. Rev. Drug. Discov. 2020, 5, 311–332. doi: 10.1038/s41573-019-0058-8

18. B. H. Gan, J. Gaynord, S. M. Rowe, T. Deingruber, D. R. Spring, Chem Soc Rev, 2021, 50, 7820. doi: 10.1039/d0cs00729c

19. K. Kornmueller, B. Lehofer, G. Leitinger, H. Amenitsch, R. Prassl Nano Res. 2018, 11 (2), 913−928. doi: 10.1007/s12274-017-1702-4

20. M.R. Yaeman, N.Y. Yount Pharmacol. Rev. 2003, 55 (1), 27−55. doi: 10.1124/pr.55.1.2

21. R.S. Christopher, A.P. Alexandra, E. Wael E, J. Dourka, J. Jury, E.F. Verdu, B. K. Coombes Nat. Commun. 2021, 12, 6664. doi: 10.1038/s41467-021-26992-4

22. R.L. Unckless, V.M. Howick, B.P. Lazzaro Curr. Biol. 2016, 26, 257–262. doi: 10.1016/j.cub.2015.11.063

23. R.L. Unckless, B.P. Lazzaro Philos. Trans. R. Soc. Lond. B. Biol. Sci. 2016, 371, 20150291. doi: 10.1098/rstb.2015.0291

24. F. Chen, B.C. Krasity, S.M. Peyer, S. Koehler, E.G. Ruby, X. Zhang, M. J McFall-Ngai mBio 2017, e00040-17. doi: 10.1128/mBio.00040-17

25. B.P. Lazzaro Curr. Opin. Microbiol., 11, 284–289. doi: 10.1016/j.mib.2008.05.001

26. M.A. Hanson, A. Dostálová, C. Ceroni, M. Poidevin, S. Kondo, B. Lemaitre ELife 2019, e44341. doi: 10.7554/elife.44341

27. R.E.W. Hancock, E.F. Haney, E.E. Gill Nat. Rev. Immunol., 16 (5), 321–334. doi: 10.1038/nri.2016.29

28. T.W. Cullen, W.B. Schofield, N.A. Barry, E. E. Putnam, E. A. Rundell, M. S. Trent Science 2015, 347, 170–175. doi: 10.1126/science.1260580

29. G. Bell, P.H. Gouyon Microbiology 2003, 149, 1367–1375. doi: 10.1099/mic.0.26265-0

30. F.J. Silva, M. Muñoz-Benavent, C. García-Ferris, A. Latorre Sci. Rep. 2020, 10, 21058. doi: 10.1038/s41598-020-77982-3

31. X.J. Pei, Y.L. Fan, Y. Bai, T. Bai, C. Schal, Z. Zhang, N. Chen, S. Li, T. Liu PLOS Biol. 2021, 19(7), e3001330. doi: 10.1371/journal.pbio.3001330

32. M.W. Zachery, E.S. Michael Sci. Rep. 2021, 11, 24196. doi: 10.1038/s41598-021-03695-w

33. J. Guzman, A. Vilcinskas Appl. Microbiol. Biotechnol. 2020, 104, 10369–10387. doi: 10.1007/s00253-020-10973-6

34. P. Das, T. Sercu, K. Wadhawan, I. Padhi, S. Gehrmann, F. Cipcigan, V. Chenthamarakshan, H. Strobelt, C. D. Santos, P. Chen, Y. Y. Yang, J. P. K. Tan, J. Hedrick, J. Crain, A. Mojsilovic Nat. Biomed. Eng. 2021, 5, 613–623. doi: 10.1038/s41551-021-00689-x

35. A. M. King, Z. Zhang, E. Glassey, P. Siuti, J. Clardy, C. A. Voigt Nat. Microbiol. 2023, 8, 2420–2434

36. D. C. Fjell, E.W.R. Hancock, H. Jenssen Curr. Pharm. Anal. 2010, 6 (2), 66−75. doi: 10.3389/fmicb.2019.03097

37. E.Y. Lee, M.W. Lee, B.M. Fulan, A. L. Ferguson, G.C. L Wong Interface. Focus. 2017, 7 (6), 20160153. doi: 10.1098/rsfs.2016.0153

38. W.F. Porto, Á.S. Pires, O.L. Franco PLoS One 2012, 7(12), No. e51444. doi: 10.1371/journal.pone.0051444

39. P. Bhadra, J. Yan, J. Li, S. Fong, S. W. I. Siu Sci. Rep. 2018, 8, 1697. doi: 10.1038/s41598-018-19752-w

40. Q. Chen, C. Yang, Y. Xie, Y. Wang, X. Li, K. Wang, J. Huang, W. Yan J. Chem. Inf. Model. 2022; 62(10): 2617-2629. doi: 10.1021/acs.jcim.2c00089

41. X. Su, J. Xu, Y. Yin, X. Quan, H. Zhang BMC. Bioinform. 2019, 20, 730. doi: 10.1186/s12859-019-3327-y

42. C.D. Fjell, J.A. Hiss, R.E.W. Hancock, G. Schneider Nat. Rev. Drug. Discov. 2012; 11 (1), 37−51. doi: 10.1038/nrd3591

43. K. Yan, H. Lv, Y. Guo, W. Peng, B. Liu Bioinformatics 2023, 39, 1, btac715. doi: 10.1093/bioinformatics/btac715

44. J. Yan, P. Bhadra, A. Li, P. Sethiya, L. Qin, H. K. Tai, K. H. Wong, S. W. Siu Mol. Ther. Nucleic. Acids. 2020, 20, 882–894. doi: 10.1016/j.omtn.2020.05.006

45. J. Huang, Y. Xu, Y. Xue, Y. Huang, X. Li, X. Chen, Y. Xu, D. Zhang, P. Zhang, J. Zhao, J. Ji Nat. Biomed. Eng. 2023, 7, 797–810. doi:10.1038/s41551-022-00991-2

46. Y. Ma, Z. Guo, B. Xia, Y. Zhang, X. Liu, Y. Yu, N. Tang, X. Tong, M. Wang, X. Ye, J. Feng, Y. Chen, J. Wang Nat. Biotechnol. 2022, 40, 921–931. doi: 10.1038/s41587-022-01226-0

47. P. Szymczak, M. Możejk, T. Grzegorzek, R. Jurczak, M. Bauer, D. Neubauer, K. Sikora, M. Michalski, J. Sroka, P. Setny, W. Kamysz, E. Szczurek Nat. Commun. 2023, 14, 1453. doi: 10.1101/2022.01.27.478054

48. Huang G, Liu Z, Laurens van der M, K. Q. Weinberger. 2017 IEEE Conference on Computer Vision and Pattern Recognition (CVPR) 2017, 2017: 2261-2269. doi: 10.1109/CVPR.2017.243

49. P. Carrasco, A.E. Pérez-Cobas, C. van de Pol, J. Baixeras, A. Moya, A. Latorre Int. Microbiol. 2014, 17, 99–109. doi: 10.2436/20.1501.01.212

50. J. Jumper, R. Evans, A. Pritzel, T. Green, M. Figurnov, O. Ronneberger, K. Tunyasuvunakool, R. Bates, A. Žídek, A. Potapenko, A. Bridgland, C. Meyer, S. A A Kohl, A. J Ballard, A. Cowie, B. Romera-Paredes, S. Nikolov, R. Jain, J. Adler, T. Back, S. Petersen, D. Reiman, E. Clancy, M. Zielinski, M. Steinegger, M. Pacholska, T. Berghammer, S. Bodenstein, D. Silver, O. Vinyals, A. W. Senior, K. Kavukcuoglu, P. Kohli, D. Hassabis Nature 2021, 596, 583–589. doi: 10.1038/s41586-021-03819-2

51. A. Datta, P. Kundu, A. Bhunia J. Colloid Interface Sci. 2016, 461, 335–345. doi: 10.1016/j.jcis.2015.09.036

52. S.P. Liu, L. Zhou, R. Lakshminarayanan, R.W. Beuerman Int. J. Pept. Res. Ther. 2010, 16, 199–213. doi: 10.1007/s10989-010-9230-z

53. Z. Dekan, S.J. Heade, M. Scanlon, B. A. Baldo, T. Lee, M. Aguilar, J. R Deuis, I. Vetter, A. G. Elliott, M. Amado, M. A Cooper, D. Alewood, P. F. Alewood Angew. Chem. 2017, 129, 8615–8619. doi: 10.1002/anie.201703360

54. W. Li, F. Lin, A. Hung, A. Barlow, M. Sani, R. Paolini, W. Singleton, J. Holden, M. A. Hossain, F. Separovic, N. M. O’Brien-Simpson, J. D. Wade Chem. Sci. 2022, 13, 2226. doi: 10.1039/d1sc05662j

55. H.T. Lee, C.C. Lee, J.R. Yang, J.Z.C. Lai, K.Y. Chang BioMed. Res. Int. 2015; 2015: 475062. doi: 10.1155/2015/475062

56. G. Wang, X. Li, Z. Wang Nucleic Acids Res. 2016; 44: D1087–D1093. doi: 10.1093/nar/gkv1278

57. U. Gawde, S. Chakraborty, F.H. Wagh, R.S. Barai, A. Khanderkar, R. Indraguru, T. Shirsat, S. Idicula-Thomas Nucleic Acids Res. 2023, 51, Database issue: D377–D383. doi: 10.1093/nar/gkac933

58. X. Zhao, H. Wu, H. Lu, G. Li, Q. Huang Plos One 2013, 8, e66557. doi: 10.1371/journal.pone.0066557

