## Supplementary material for "The discovery of antimicrobial peptides from the gut microbiome of cockroach *Blattella germanica* using deep learning pipeline": All supplementary materials and data: Cover Letter.docx

***Short running title*:** Antimicrobial peptides from cockroach gut microbiome.

*What does the current manuscript add to the existing knowledge? (Research Highlights)*

1- Proposed a new efficient computational tool (AMPidentifer) for the discovery of antimicrobial peptides (AMPs).
2- Identified AMPs from the gut microbiome of cockroach, *Blattella germanica* for the first time.
3- Experimentally verified two new AMPs with potent antimicrobial activities and low mammalian cytotoxicity.
4- Explored antimicrobial influences of single Cys in identified AMPs, and contributed insights to future rational design.
5- One of the newly identified AMPs showed good wound healing effects and antimicrobial activities in vivo.
6- Demonstrated the feasibility of the interdisciplinary strategy for identifying biocompatible AMPs from nature.

**Provide any other information below:**

We would like to submit the manuscript entitled: **The discovery of antimicrobial peptides from the gut microbiome of cockroach *Blattella germanica* using deep learning.** We affirm that the submission represents original work that has not been published previously and is not currently being considered or submitted to another journal, until a decision has been made. Also, we confirm that each author has seen and approved the contents of the submitted manuscript. The experimental procedures involving mice was approved by the Health Science Center of Xi'an Jiaotong University (New Permit Number: XJTUAE2023-2129).

Tian Liu

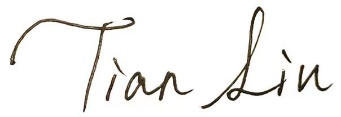

School of Bioengineering, Dalian University of Technology, Dalian 116024
Tel. Number: 86-411-84706527
