## Supplementary material for "The discovery of antimicrobial peptides from the gut microbiome of cockroach *Blattella germanica* using deep learning pipeline": All supplementary materials and data: Manuscript.docx

$Precision=\frac{TP}{TP+FP}$	(13)

$Sensitivity=\frac{TP}{TP+FN}$	(14)

$Specificity=\frac{TN}{TN+FP}$	(15)

Where the TP, FP, TN, and FN indicated true positive number, false positive number, true negative number, and false negative number, respectively.

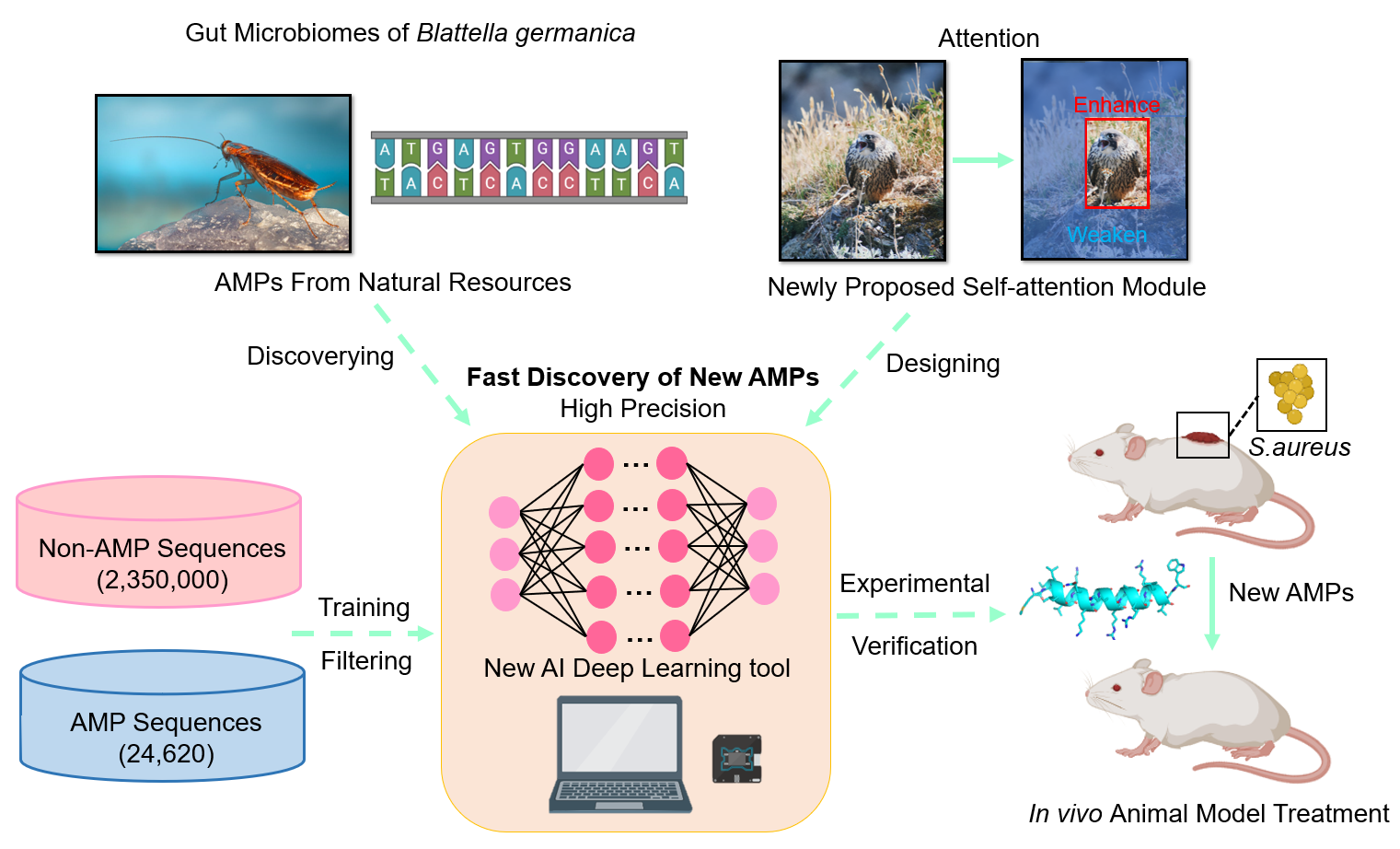
 **Figure Legends**

**The Graphic Abstract**

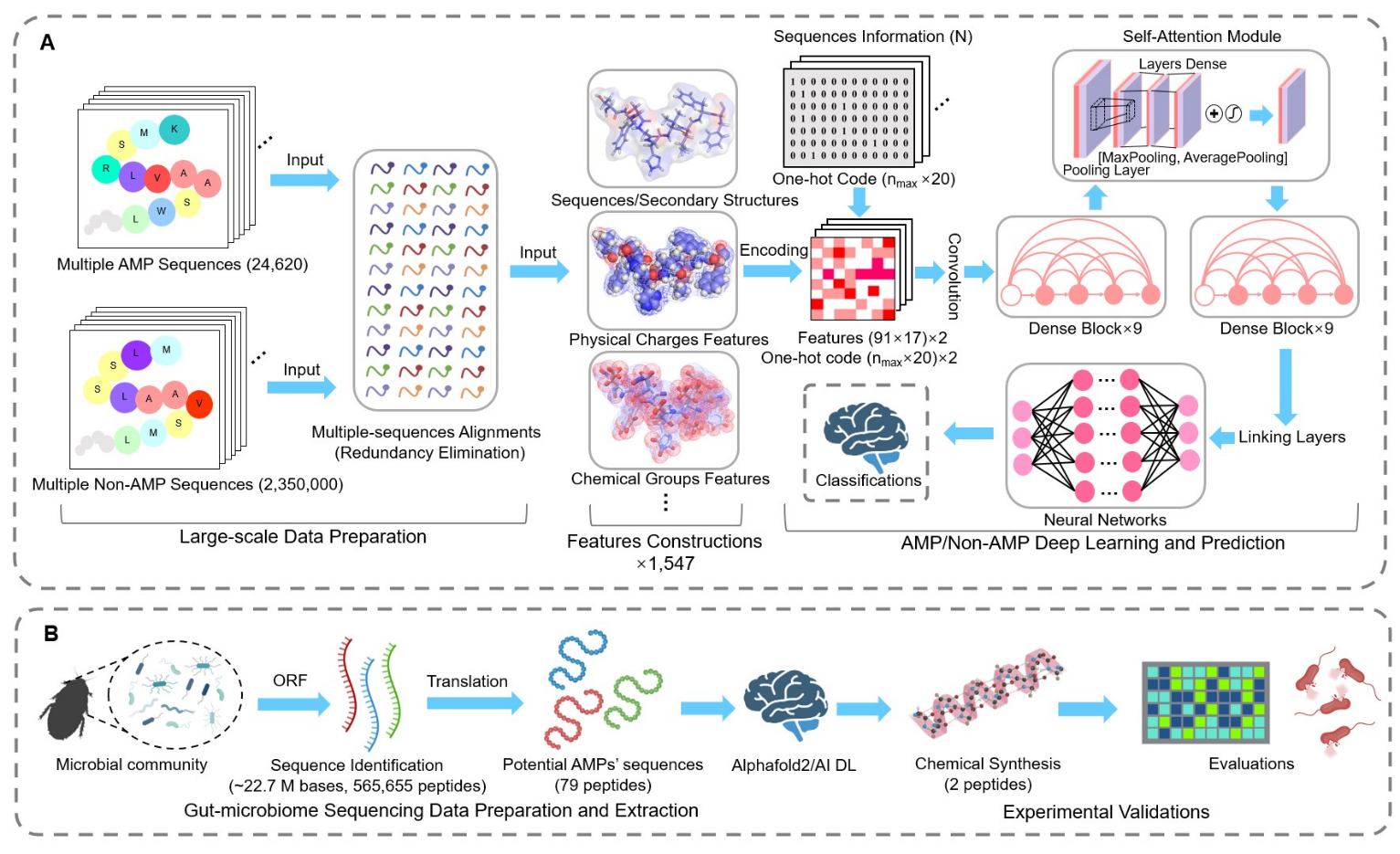

**Fig. 1. Schematic representation of AMPidentifer workflow.** We mined gut microbiome data for potential AMPs, resulting in candidate AMPs for chemical synthesis and in vitro validations. **A**, Deep learning algorithms for identifying AMPs and non-AMPs, in which multiple features including physical, chemical, sequences, and structural information are utilized to construct input information for the model. **B**, AMPs mining strategies from gut microbiome data collected from pathogenic insect *B. germanica*.

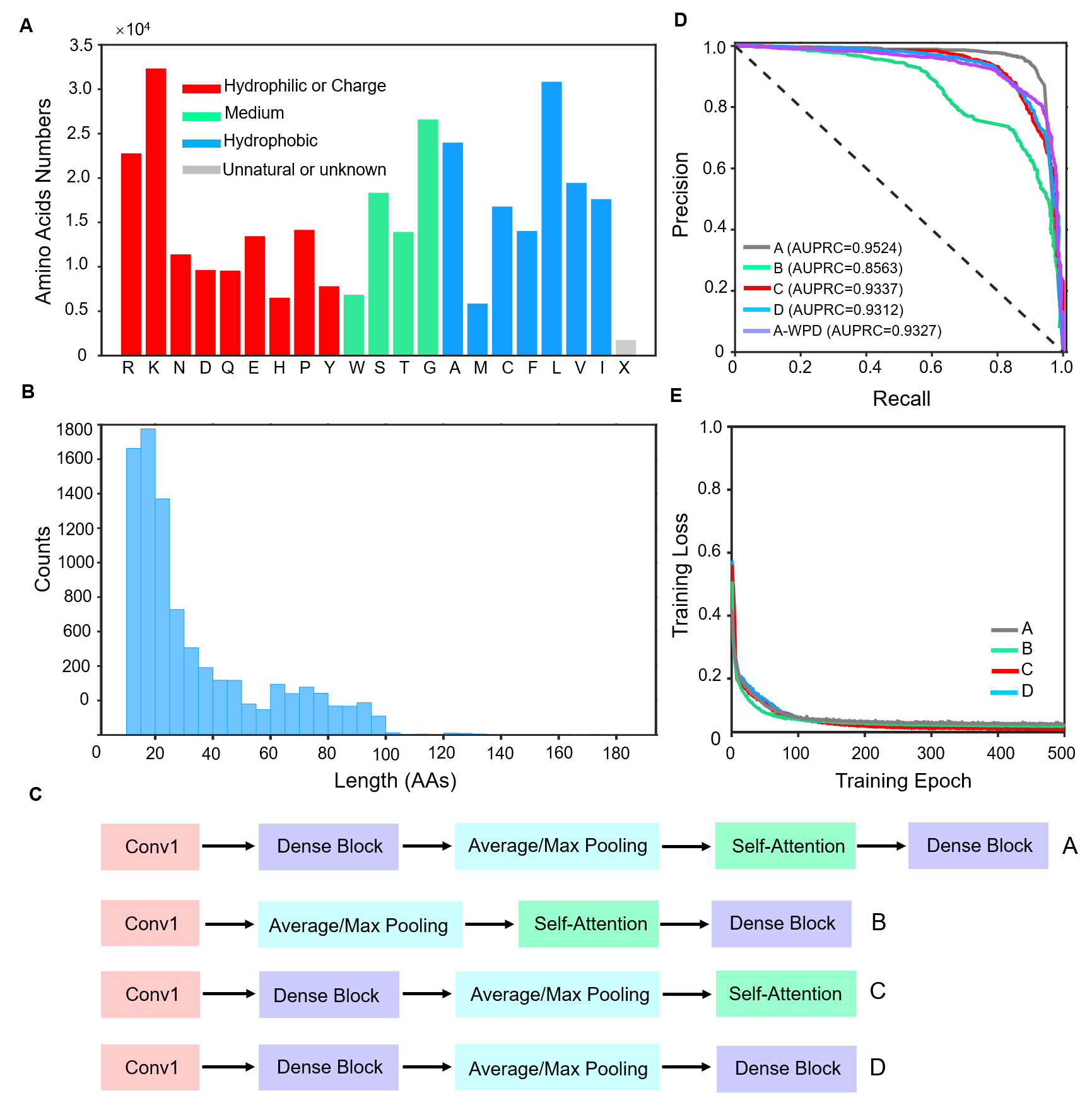

**Fig. 2. Establishing AMP prediction deep learning models.** **A**, Amino acids distribution of AMPs collected from various databases for model training. X represents unknown or unconventional amino acids. **B**, the Length distribution of AMPs collected from various databases. **C**, Summary of four model strategies for testing and building. **(D)**, AUPRC evaluations for different combinations of strategies A, B, C, D, and A-without protein descriptors (A-WPD). **(E)**, Training Loss and circulation periods for different combinations strategies.

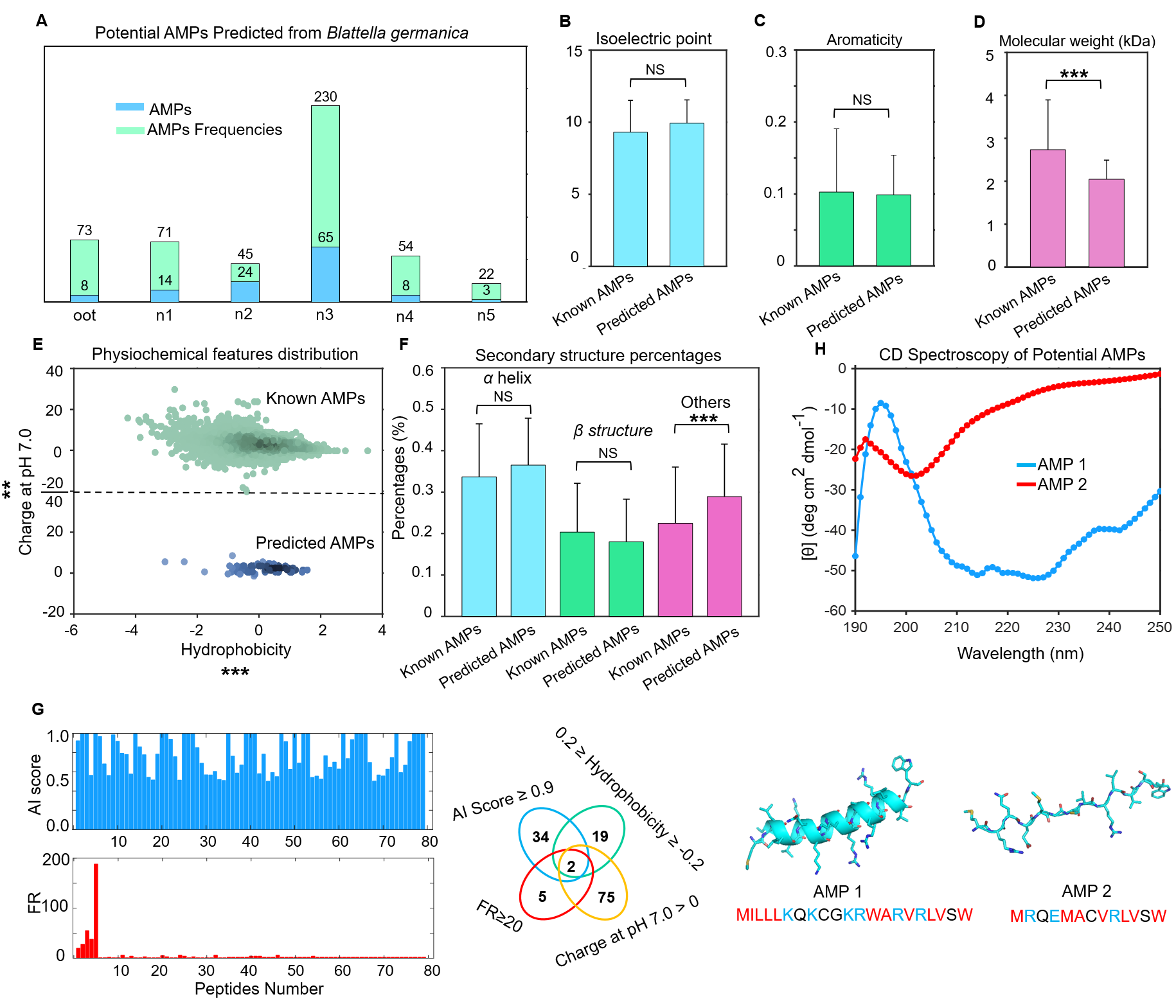

**Fig. 3. Mining potential AMPs from the gut microbiome of** ***B. germanica*. A**, The statistics of potential AMP sequences predicted by the AI DL model from different growth stages (oot, n1, n2, n3, n4, and n5) of the gut microbiome of *B. germanica*. **B-F**, The comparison of physiochemical features of isoelectric points, aromaticity, molecular weight, hydrophobicity, charge at pH 7.0, and the secondary structure (*α* helix, *β* structure, and others) percentages between known AMPs and AI-predicted AMPs. **G**, the AI score and frequencies of 79 potential AMPs from *B. germanica*. Two sequences possessing AI scores> 0.9 and frequencies > 5, were further manually selected according to their sequential features for experimental validation. The structure of two peptide structures predicted by Alphafold2. **H**, The CD spectroscopy of secondary structures of two potential AMPs with concentrations of 0.2 mg/mL in deionized water.

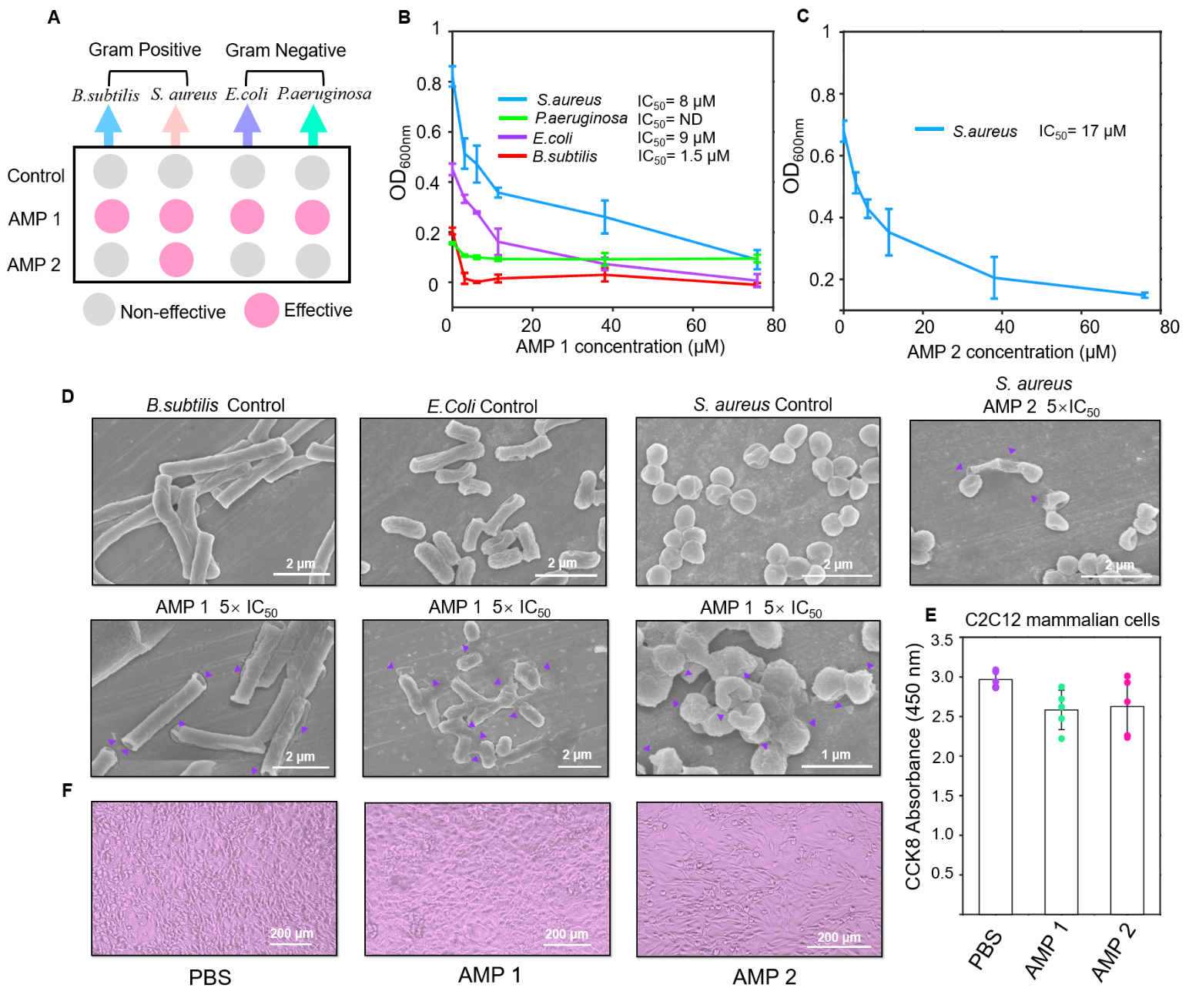

**Fig. 4. Experiments verify the strong potencies of functional AMPs.** **A**, The screen of bacterial inhibition of four peptides against *S. aureus*, *B. subtilis*, *E. coli DH5α,* and *P. aeruginosa* at the concentration of 200 μM. **B**, The corresponding IC_50_ values and inhibition effects of AMP 1 and **C,** AMP 2 towards different bacteria at concentrations of 0 μM, 3.8 μM, 6 μM, 11 μM, 38 μM, and 76 μM cultivating for 7-8 h, respectively. **D,** The SEM examination of *Staphylococcus aureus*, *Bacillus subtilis*, and *Escherichia coli DH5α* cells treated with AMPs, showed cell content leakage and disruption of cell wall/membrane. All ruptured cells were indicated by a purple arrow. **E,** The C2C12 mammalian cells with 0.5% PBS solution or 100 μM AMPs were cultivated for 15 h at 37 °C. The cell activities and survival rates were indicated by the absorbance of CCK8 at 450 nm with 5 independent replicates (average survival rates＝87±8.4% and 88.5±12.3%, respectively). **F,** the representative optical microscopy images of C2C12 cells.

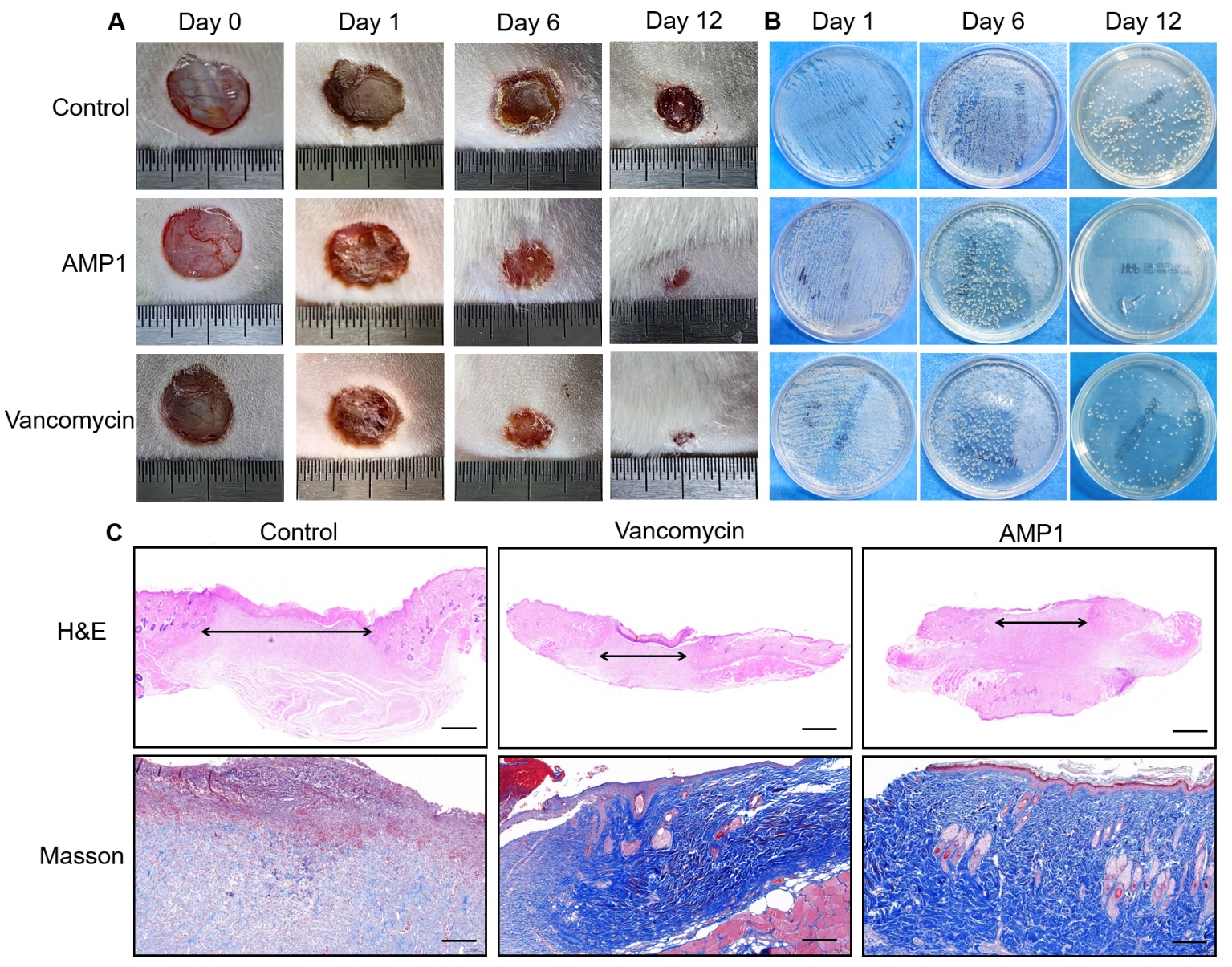

**Fig. 5. The antimicrobial and wound healing therapeutic effects of AMP 1 *in vivo*. A,** the representative wound healing results of the control PBS, Vancomycin, and AMP 1 to infected mice model at the same dosage *in vivo*. **B,** The experimental verifications of infected *S.aureus* load of the control group, antibiotic Vancomycin treatment group, and AMP 1 treatment group by streaking on LB agar medium plate. **C,** The representative histological images of infection tissues from the control group, Vancomycin treatment group, and AMP 1 treatment group generated by experimental H&E and Masson dye methods

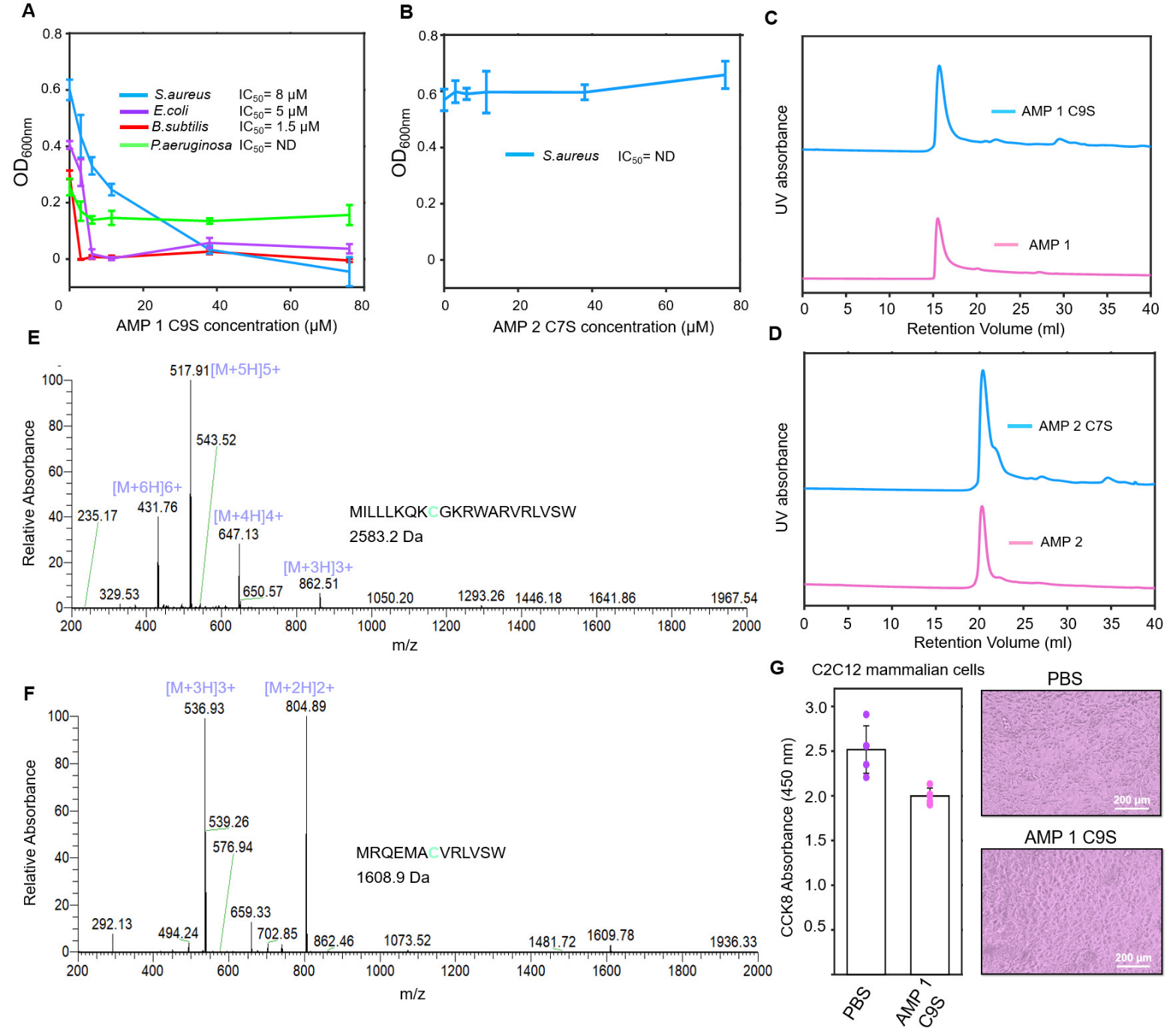

**Fig. 6.** **The influences of Cys residues in antimicrobial activities. A,** AMP 1 C9S and **B,** AMP 2 C7S towards different pathogenic bacteria at concentrations of 0, 3.8, 6, 11, 38, and 76 μM at 37 °C cultivating for 7-8 h, respectively. The Mass spectrometry of **C,** AMP 1 and **D,** AMP 2 diluted in deionized water at the concentrations of 2 mg/mL. The SECresults of **E,** AMP 1and AMP 1 C9S, **F,** AMP 2and AMP 2 C7S at 25 °C in buffer containing 150 mM NaCl, 20 mM Tris, pH 7.0. **G,** The C2C12 mammalian cells with 0.5% PBS solution or 100 μM AMP 1 C9S were cultivated for 15 h at 37 °C. The cell activities and survival rates were indicated by the absorbance of CCK8 at 450 nm with 5 independent replicates (average survival rates＝80±3.5%).

**Table 1.** **Comparison of the highest performance of different model designing strategies**

| **Method** | **TP** | **FN** | **TN** | **FP** | **Sensitivity (%)** | **Specificity (%)** | **Precision (%)** | **AUPRC** |
| --- | --- | --- | --- | --- | --- | --- | --- | --- |
| A | 849 | 141 | 38,207 | 56 | 85.75 | 99.85 | 93.81 | 0.9524 |
| B | 892 | 98 | 37,970 | 293 | 90.01 | 99.23 | 75.27 | 0.8563 |
| C | 901 | 89 | 38,064 | 199 | 91.01 | 99.48 | 77.94 | 0.9337 |
| D | 928 | 62 | 38,008 | 255 | 93.74 | 99.33 | 78.44 | 0.9312 |

**Table 2.** **Comparison of the highest performance among frequently-used AMPs prediction algorithms ^8-9, 35-46^**

| **Method** | **TP** | **FN** | **TN** | **FP** | **Sensitivity (%)** | **Specificity (%)** | **Precision (%)** | **AUPRC** |
| --- | --- | --- | --- | --- | --- | --- | --- | --- |
| Our Pipeline | 849 | 141 | 38,207 | 56 | 85.75 | 99.85 | 93.81 | 0.9524 |
| Random Forest | 840 | 150 | 38,140 | 123 | 84.84 | 99.69 | 87.23 | 0.9226 |
| XGBoost | 893 | 97 | 38,154 | 89 | 90.20 | 99.77 | 90.93 | 0.9498 |
| Decision Tree | 863 | 127 | 37,592 | 671 | 87.17 | 98.25 | 56.26 | 0.9271 |
| Linear Regression | 469 | 521 | 38,093 | 170 | 47.37 | 99.56 | 73.40 | 0.6083 |
| Dense-Net (CNN) | 825 | 165 | 38,080 | 183 | 83.33 | 99.52 | 81.85 | 0.8762 |
| LSTM | 766 | 224 | 38,117 | 146 | 77.37 | 99.62 | 83.99 | 0.8856 |
| Transformer | 739 | 251 | 38,023 | 240 | 74.65 | 99.37 | 75.49 | 0.8212 |

**Table 3. Antimicrobial activities of the identified AMPs and mutated sequences**

| **AMP** | **Bacteria Species** | **IC_50_ (μM)** | **MIC (μM)** |
| --- | --- | --- | --- |
| AMP 1 | *B. subtili* | 1.5 | 6.0 |
|  | *S. aureus* | 8.0 | 85.0 |
|  | *E. Coli* | 9.0 | 45.0 |
|  | *P. aeruginosa* | ND | ND |
|  | *B. subtili* | ND | ND |
| AMP 2 | *S. aureus* | 17.0 | 92.5 |
|  | *E. Coli* | ND | ND |
|  | *P. aeruginosa* | ND | ND |
|  | *B. subtili* | 1.5 | 6.0 |
| AMP 1 C9S | *S. aureus* | 8.0 | 38.0 |
|  | *E. coli* | 5.0 | 7.0 |
|  | *P. aeruginosa* | ND | ND |
|  | *B. subtili* | ND | ND |
| AMP 2 C7S | *S. aureus* | ND | ND |
|  | *E. coli* | ND | ND |
|  | *P. aeruginosa* | ND | ND |

**Conflict of Interest:** The authors declare that they have no known competing financial interests or personal relationships that could have appeared to influence the work reported in this paper.

**Reference**

**[1]** E.Y. Klein, M. Milkowska-Shibata, K.K. Tseng, M. Sharland, S. Gandra, C. Pulcini, R. Laxminarayan *Lancet. Infect. Dis.* **2021**, *21*, 107–115. doi: 10.1016/S1473-3099(20)30332-7

**[22]** R.L. Unckless, V.M. Howick, B.P. Lazzaro *Curr. Biol.* **2016**, *26*, 257–262. doi: https://doi.org/10.1016/j.cub.2015.11.063

**[23]** R.L. Unckless, B.P. Lazzaro *Philos. Trans. R. Soc. Lond. B. Biol. Sci.* **2016**, *371*, 20150291. doi: 10.1098/rstb.2015.0291

**[24]** F. Chen, B.C. Krasity, S.M. Peyer, S. Koehler, E.G. Ruby, X. Zhang, M. J McFall-Ngai *mBio* **2017**, *8*, e00040-17. doi: 10.1128/mBio.00040-17

**[25]** B.P. Lazzaro *Curr. Opin. Microbiol.* **2008**, *11*, 284–289. doi: 10.1016/j.mib.2008.05.001

**[26]** M.A. Hanson, A. Dostálová, C. Ceroni, M. Poidevin, S. Kondo, B. Lemaitre *ELife* **2019**, *8*, e44341. doi: 10.7554/elife.44341

**[27]** R.E.W. Hancock, E.F. Haney, E.E. Gill *Nat. Rev. Immunol.* **2016**, *16 (5)*, 321–334. doi: 10.1038/nri.2016.29
