## Supplementary material for "The discovery of antimicrobial peptides from the gut microbiome of cockroach *Blattella germanica* using deep learning pipeline": All supplementary materials and data: Manuscript.pdf

**S Chen[a,d,#], H Qi[a,#], X Zhu[c,#], T Liu[b], Y Teng[a], Q Gong[c,*], C Jia[b,*], T Liu[a,*]**
[a]*MOE Key Laboratory of Bio-Intelligent Manufacturing, School of Bioengineering, Dalian University of Technology, Dalian, Liaoning, 116024, China.*
[b]*School of Science, Dalian Maritime University, 116026, China.*
[c]*The First Affiliated Hospital of Xi'an Jiaotong University, Xian 710061, China.*
[d]*Division of Biomedical Science, Chinese University of Hong Kong, Shatin, NT, Hong Kong SAR, China.*

**[#]Co-First Authors, these authors contributed equally to this study**
***Corresponding authors**

***Corresponding Author:**

$$Precision = \frac{TP}{TP+FP} \quad (13)$$

$$Sensitivity = \frac{TP}{TP+FN} \quad (14)$$

$$Specificity = \frac{TN}{TN+FP} \quad (15)$$

Where the TP, FP, TN, and FN indicated true positive number, false positive number, true negative number, and false negative number, respectively.

### Figure Legends

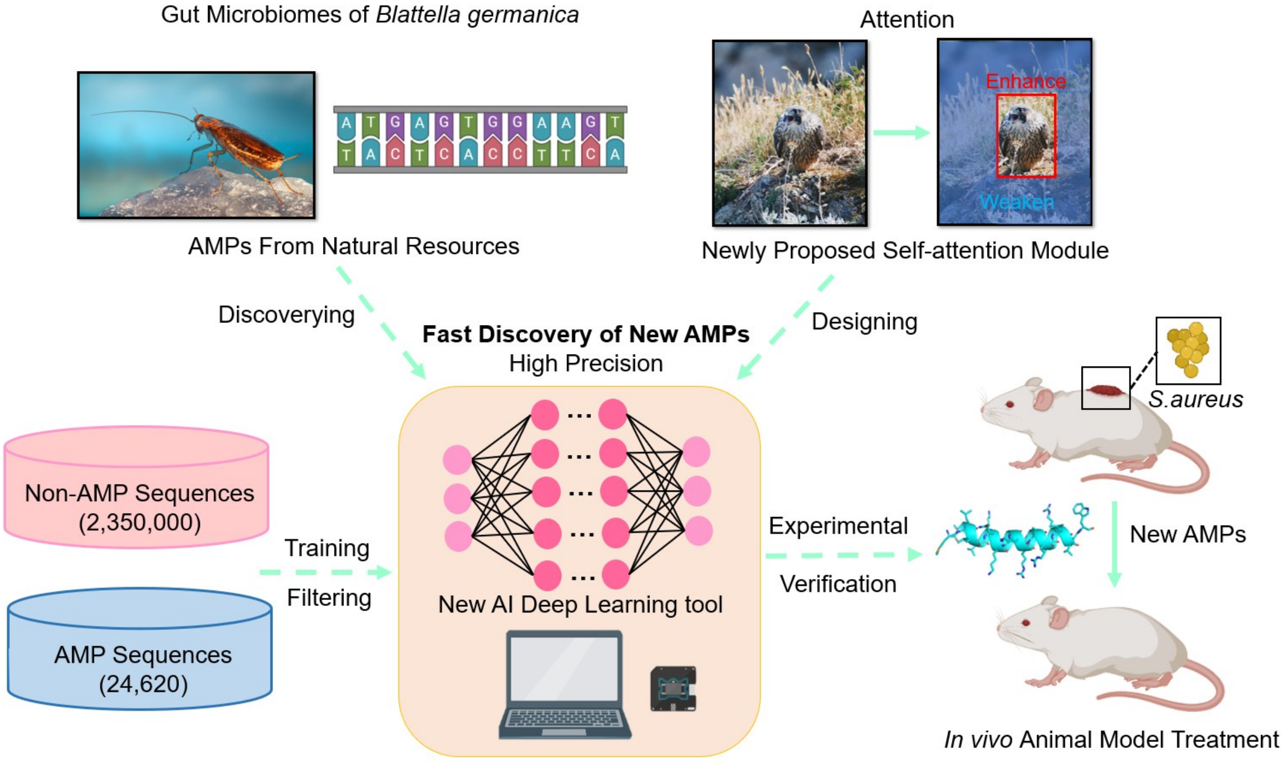


**The Graphic Abstract**

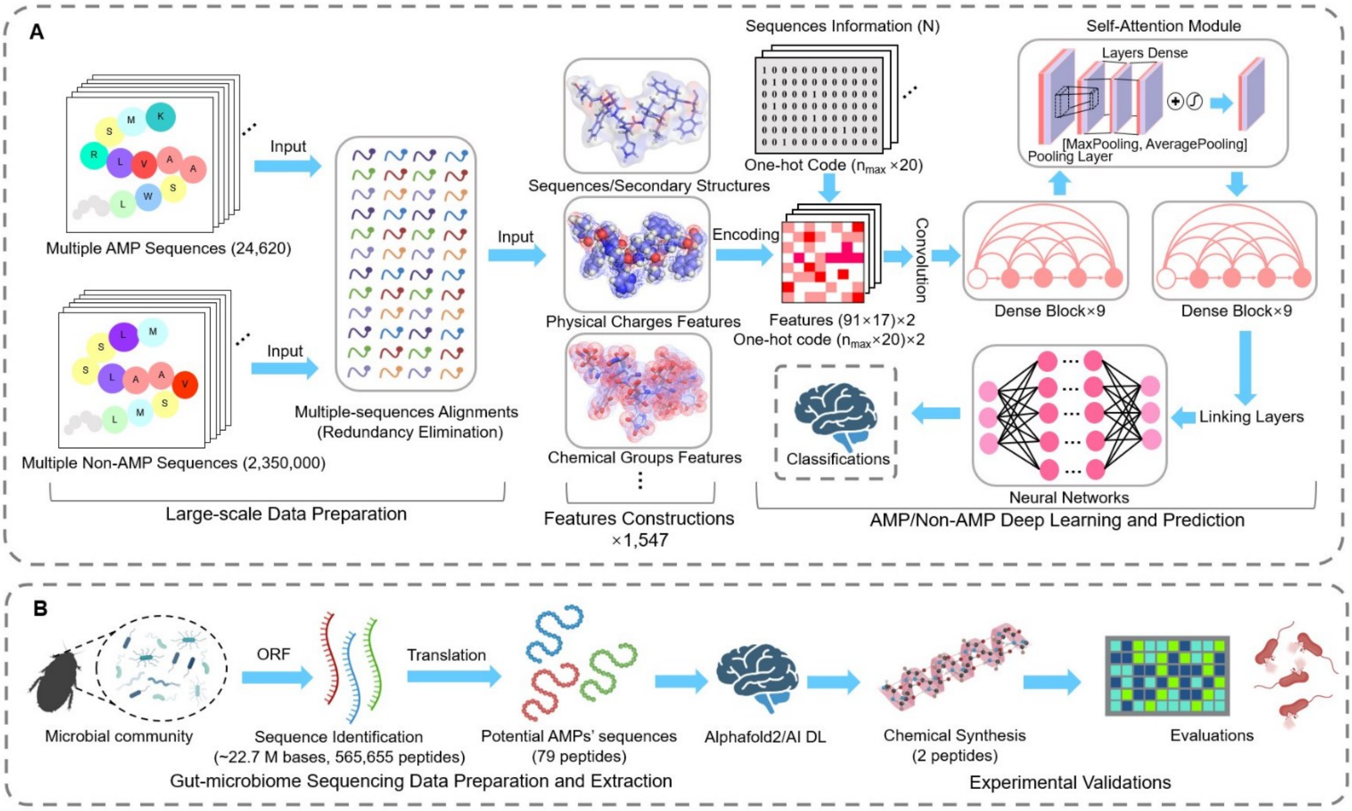


**Fig. 1. Schematic representation of AMPidentifer workflow.** We mined gut microbiome data for potential AMPs, resulting in candidate AMPs for chemical synthesis and in vitro validations. **A**, Deep learning algorithms for identifying AMPs and non-AMPs, in which multiple features including physical, chemical, sequences, and structural information are utilized to construct input information for the model. **B**, AMPs mining strategies from gut microbiome data collected from pathogenic insect *B. germanica*.

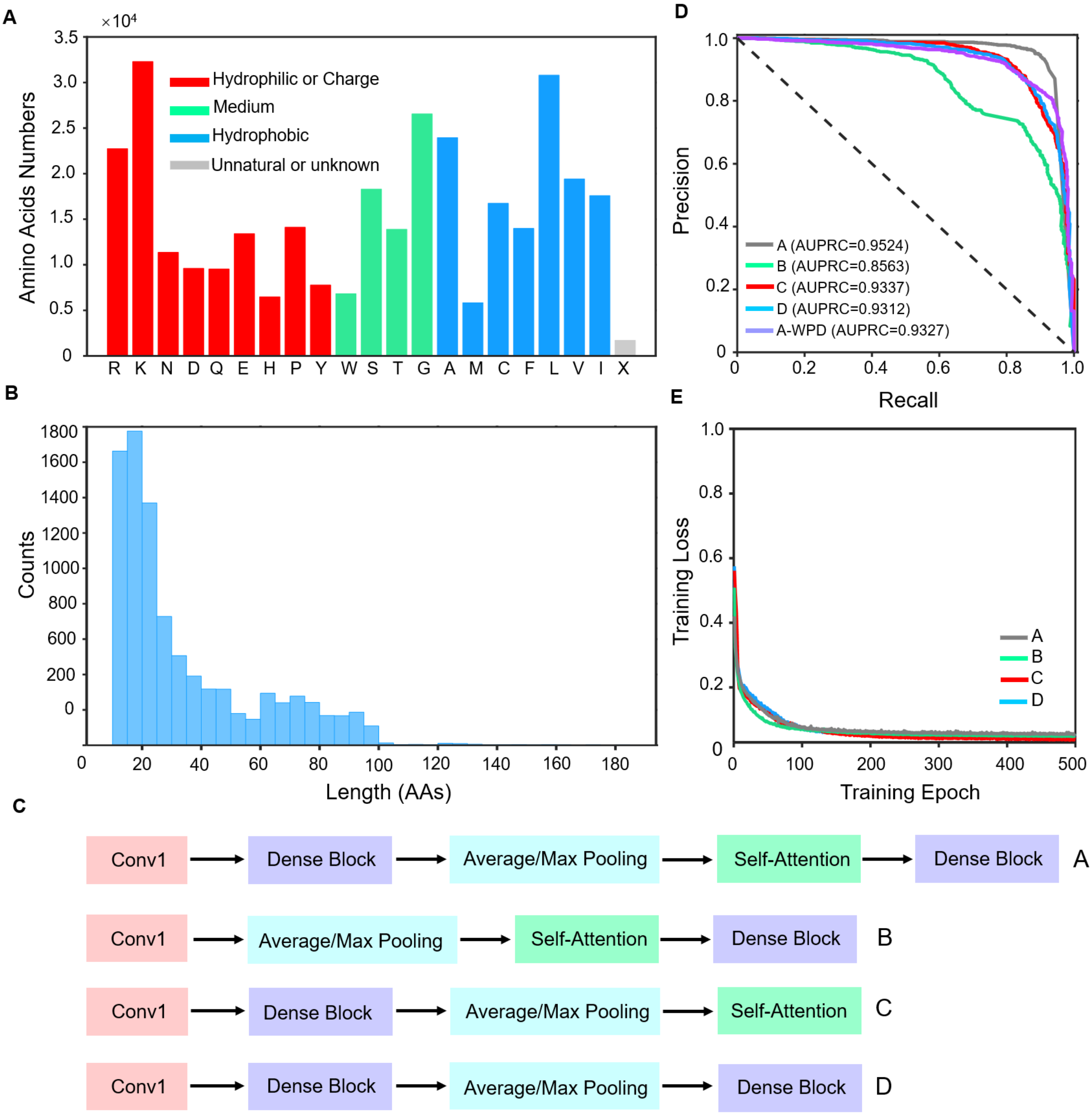


**Fig. 2. Establishing AMP prediction deep learning models. A**, Amino acids distribution of AMPs collected from various databases for model training. X represents unknown or unconventional amino acids. **B**, the Length distribution of AMPs collected from various databases. **C**, Summary of four model strategies for testing and building. **(D)**, AUPRC evaluations for different combinations of strategies A, B, C, D, and A-without protein descriptors (A-WPD). **(E)**, Training Loss and circulation periods for different combinations strategies.

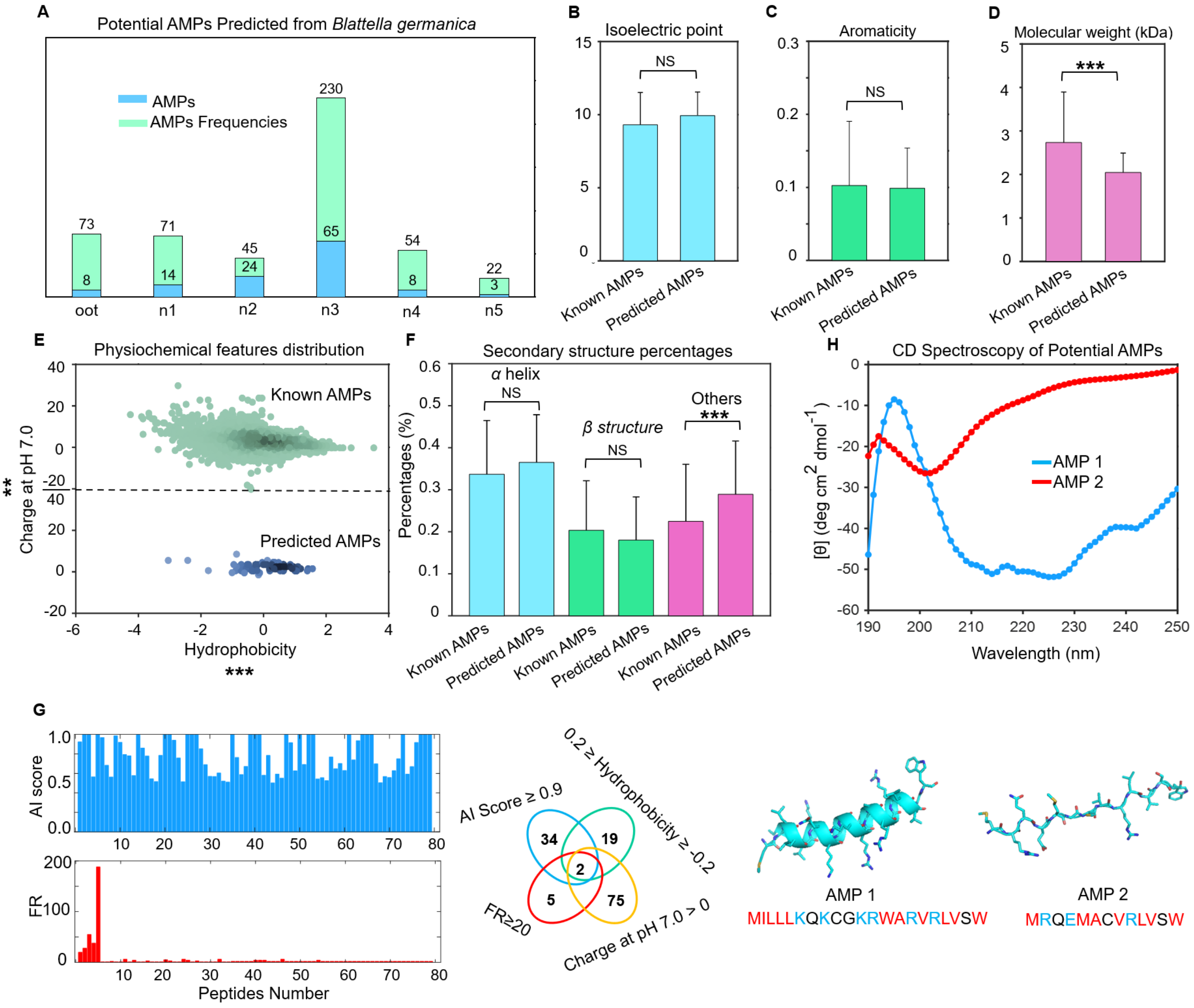


**Fig. 3. Mining potential AMPs from the gut microbiome of *B. germanica*.** **A**, The statistics of potential AMP sequences predicted by the AI DL model from different growth stages (oot, n1, n2, n3, n4, and n5) of the gut microbiome of *B. germanica*. **B-F**, The comparison of physiochemical features of isoelectric points, aromaticity, molecular weight, hydrophobicity, charge at pH 7.0, and the secondary structure (*α* helix, *β* structure, and others) percentages between known AMPs and AI-predicted AMPs. **G**, the AI score and frequencies of 79 potential AMPs from *B. germanica*. Two sequences possessing AI scores> 0.9 and frequencies > 5, were further manually selected according to their sequential features for experimental validation. The structure of two peptide structures predicted by Alphafold2. **H**, The CD spectroscopy of secondary structures of two potential AMPs with concentrations of 0.2 mg/mL in deionized water.

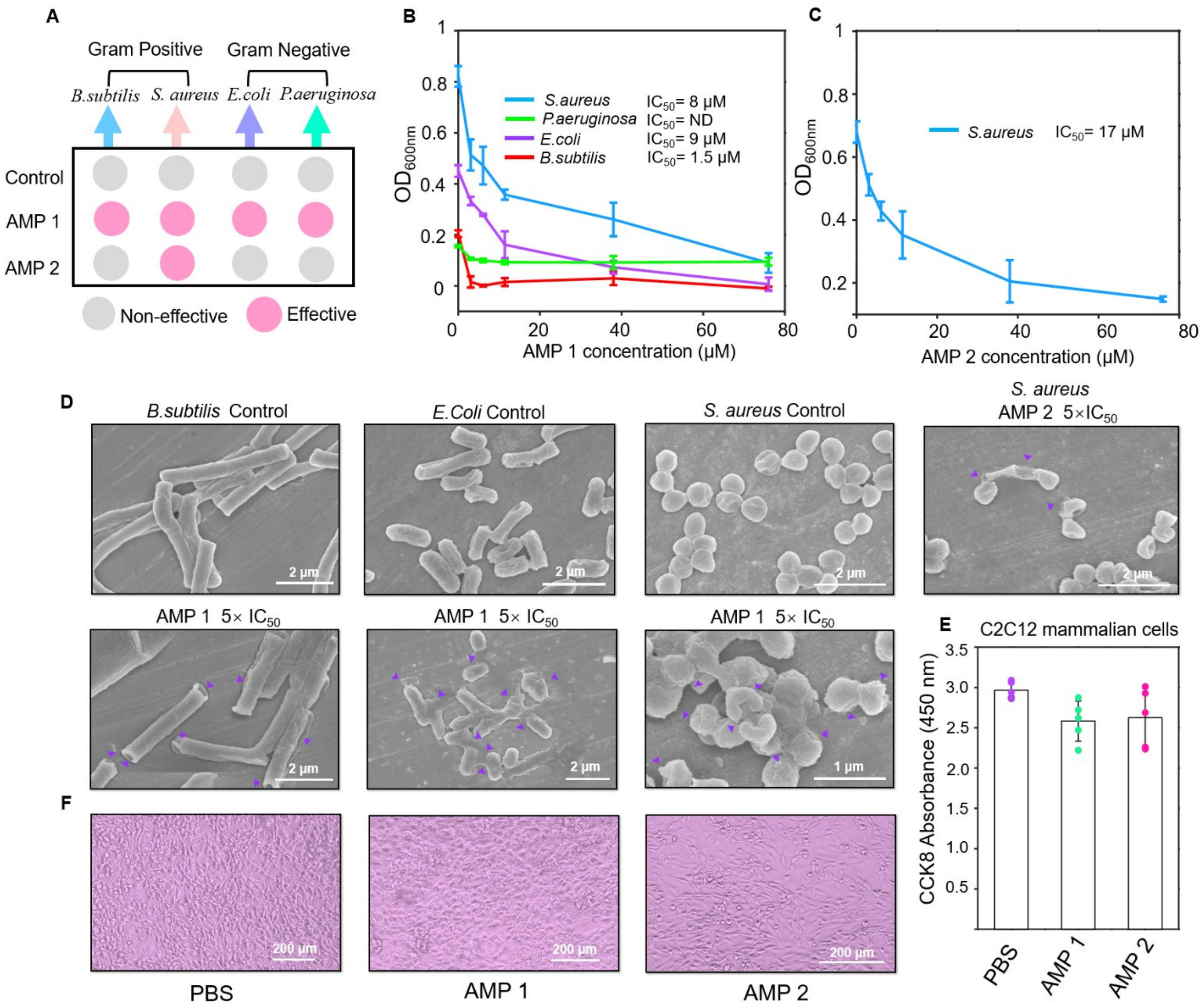


**Fig. 4. Experiments verify the strong potencies of functional AMPs.** **A**, The screen of bacterial inhibition of four peptides against *S. aureus*, *B. subtilis*, *E. coli DH5α,* and *P. aeruginosa* at the concentration of 200 µM. **B**, The corresponding $IC_{50}$ values and inhibition effects of AMP 1 and **C,** AMP 2 towards different bacteria at concentrations of 0 µM, 3.8 µM, 6 µM, 11 µM, 38 µM, and 76 µM cultivating for 7-8 h, respectively. **D,** The SEM examination of *Staphylococcus aureus*, *Bacillus subtilis*, and *Escherichia coli DH5α* cells treated with AMPs, showed cell content leakage and disruption of cell wall/membrane. All ruptured cells were indicated by a purple arrow. **E,** The C2C12 mammalian cells with 0.5% PBS solution or 100 µM AMPs were cultivated for 15 h at 37 °C. The cell activities and survival rates were indicated by the absorbance of CCK8 at 450 nm with 5 independent replicates (average survival rates = 87±8.4% and 88.5±12.3%, respectively). **F,** the representative optical microscopy images of C2C12 cells.

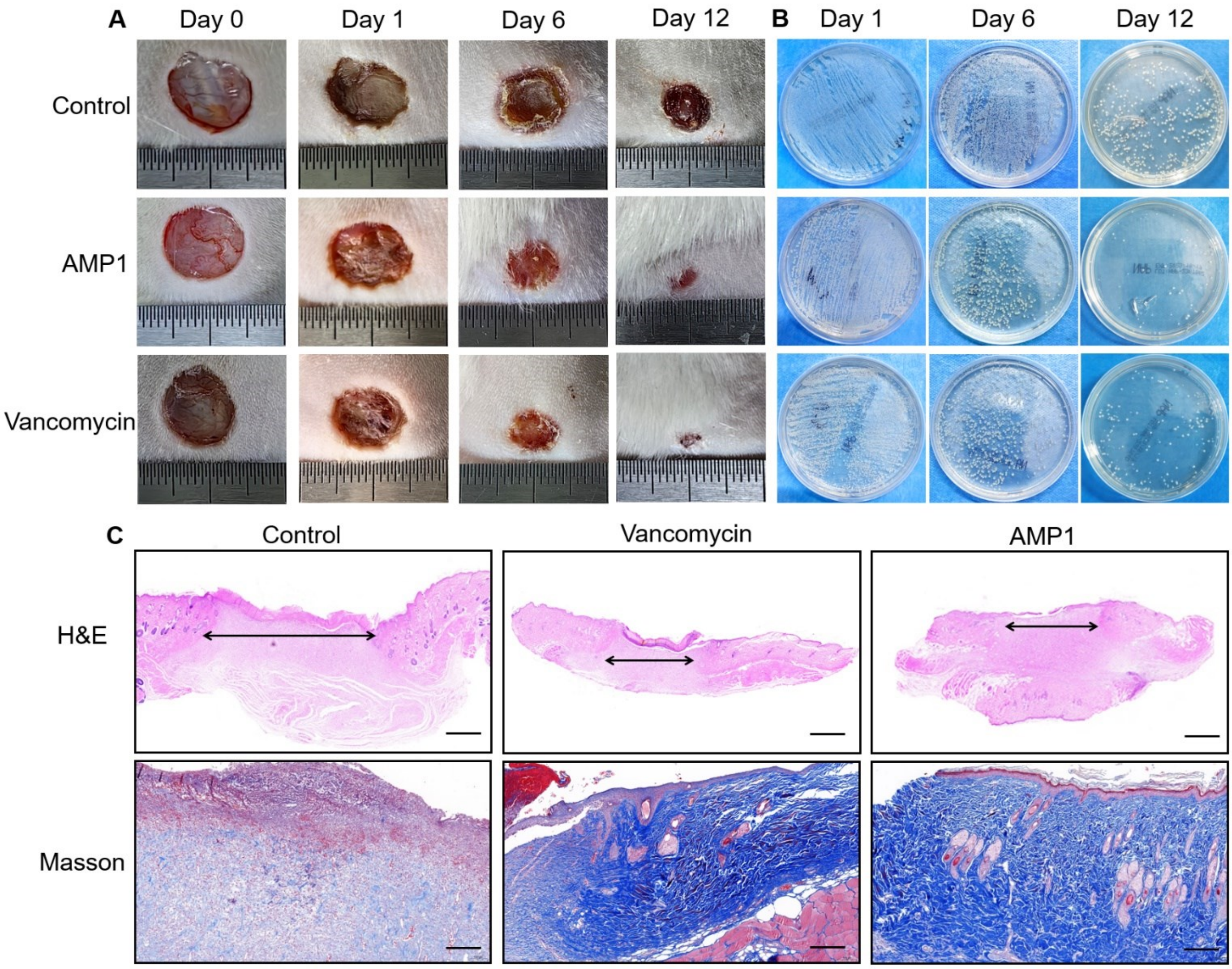


443 **Fig. 5. The antimicrobial and wound healing therapeutic effects of AMP 1 *in vivo*. A,** the representative wound
444 healing results of the control PBS, Vancomycin, and AMP 1 to infected mice model at the same dosage *in vivo*. **B,** The
445 experimental verifications of infected *S.aureus* load of the control group, antibiotic Vancomycin treatment group, and
446 AMP 1 treatment group by streaking on LB agar medium plate. **C,** The representative histological images of infection
447 tissues from the control group, Vancomycin treatment group, and AMP 1 treatment group generated by experimental
448 H&E and Masson dye methods

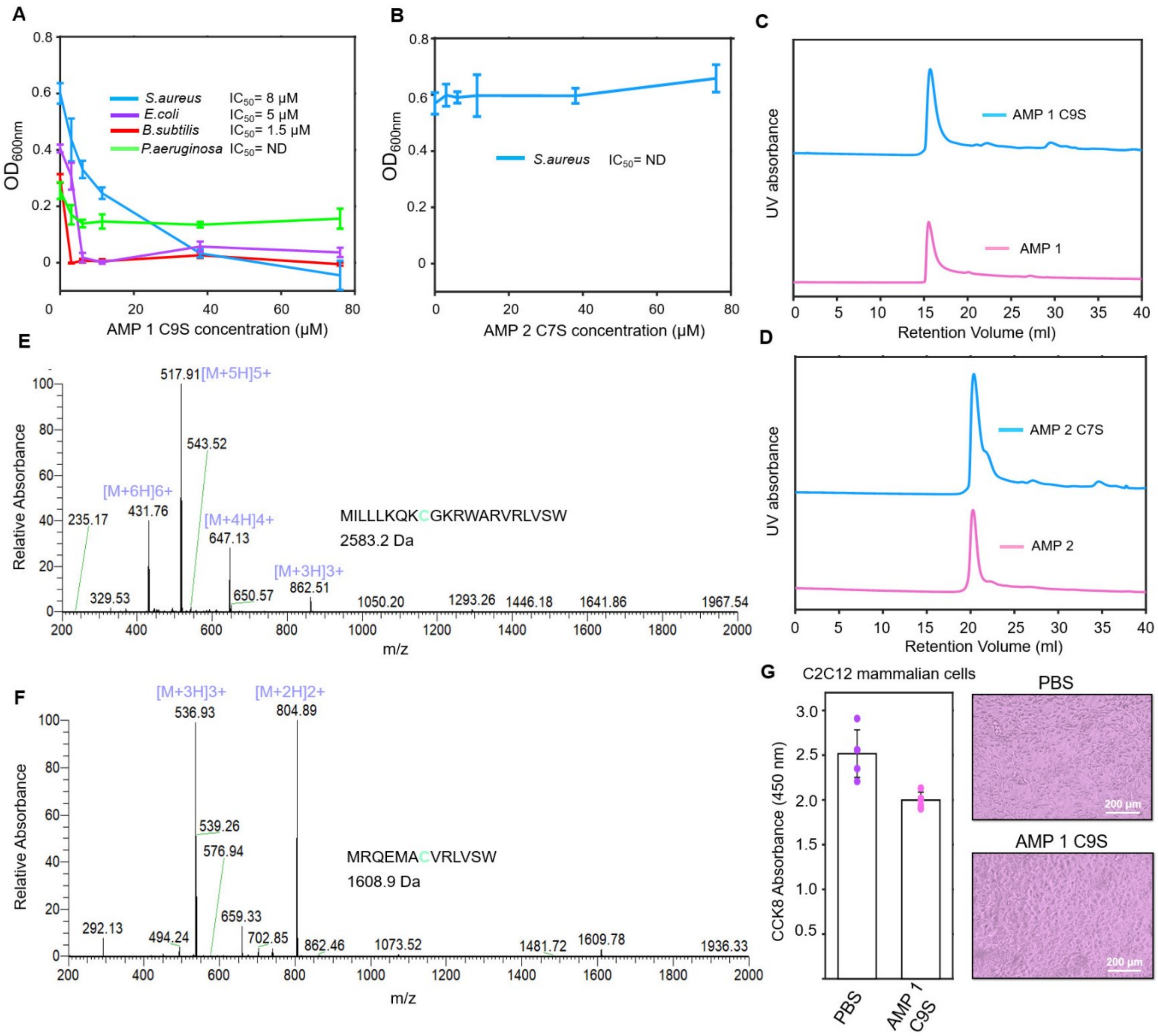


**Fig. 6. The influences of Cys residues in antimicrobial activities. A,** AMP 1 C9S and **B,** AMP 2 C7S towards different pathogenic bacteria at concentrations of 0, 3.8, 6, 11, 38, and 76 μM at 37 °C cultivating for 7-8 h, respectively. The Mass spectrometry of **C,** AMP 1 and **D,** AMP 2 diluted in deionized water at the concentrations of 2 mg/mL. The SECresults of **E,** AMP 1and AMP 1 C9S, **F,** AMP 2and AMP 2 C7S at 25 °C in buffer containing 150 mM NaCl, 20 mM Tris, pH 7.0. **G,** The C2C12 mammalian cells with 0.5% PBS solution or 100 μM AMP 1 C9S were cultivated for 15 h at 37 °C. The cell activities and survival rates were indicated by the absorbance of CCK8 at 450 nm with 5 independent replicates (average survival rates = 80±3.5%).
