## Supplementary material for "The discovery of antimicrobial peptides from the gut microbiome of cockroach *Blattella germanica* using deep learning pipeline": All supplementary materials and data: Supplementary Materials.docx

**New Self-attention module design**

Here we designed a new self-attention module, which was proposed for the first time and was demonstrated to be more effective than a 9-layers dense block in this work. In the self-attention module, the input matrix was applied with average pooling and max pooling strategy, and the outputs were concatenated together.

$F_{kij}={max}_{(p,q)\in R_{ij}}x_{kpq}$	(1)

Where *F_kij_* represents the max pooling output of the rectangular region *R_ij_* in the *k^th^* input matrix, and *x_kpq_* represents the element distributing in (p,q) position of region *R_ij_*.

$D_{kij}={average}_{(p,q)\in R_{ij}}x_{kpq}$	(2)

Where *D_kij_* represents the average pooling output of the rectangular region *R_ij_* in the *k^th^* input matrix, and *x_kpq_* represents the element distributing in (p,q) position of region *R_ij_*.

$A_{2k}=\mathrm{concatenate}\left( F_{k}, D_{k} \right)$	(3)

Where *A_2k_* represents the concatenated matrixes. Then, the channel sizes of *F_k_* and *D_k_* matrixes were decreased to s by the Keras Dense1 function, and then activated by the ReLU function.

$H_{s}=D\mathrm{ense}1(F_{k})$	(4)

$G_{s}=D\mathrm{ense}1(D_{k})$	(5)

$Q_{s}=\mathrm{ReLU}(H_{s})$	(6)

$J_{s}=\mathrm{ReLU}(G_{s})$	(7)

After this, the channel sizes of outputs were then increased to 2k by the Keras Dense2 function, and matrixes were added together.

${P1}_{2k}=D\mathrm{ense}2(Q_{s})$	(8)

${P2}_{2k}=D\mathrm{ense}2(J_{s})$	(9)

$P_{2kij}={P1}_{2kij}+{P2}_{2kij}$	(10)

In the end, *U_kij_* was activated by sigmoid function and multiplied to initial output matrixes *A_2kij_* obtained in average pooling/max pooling steps.

$U_{2k}=\mathrm{Sigmoid}(P_{2k})$	(11)

$I_{2kij}=A_{2kij} U_{2kij}$	(12)

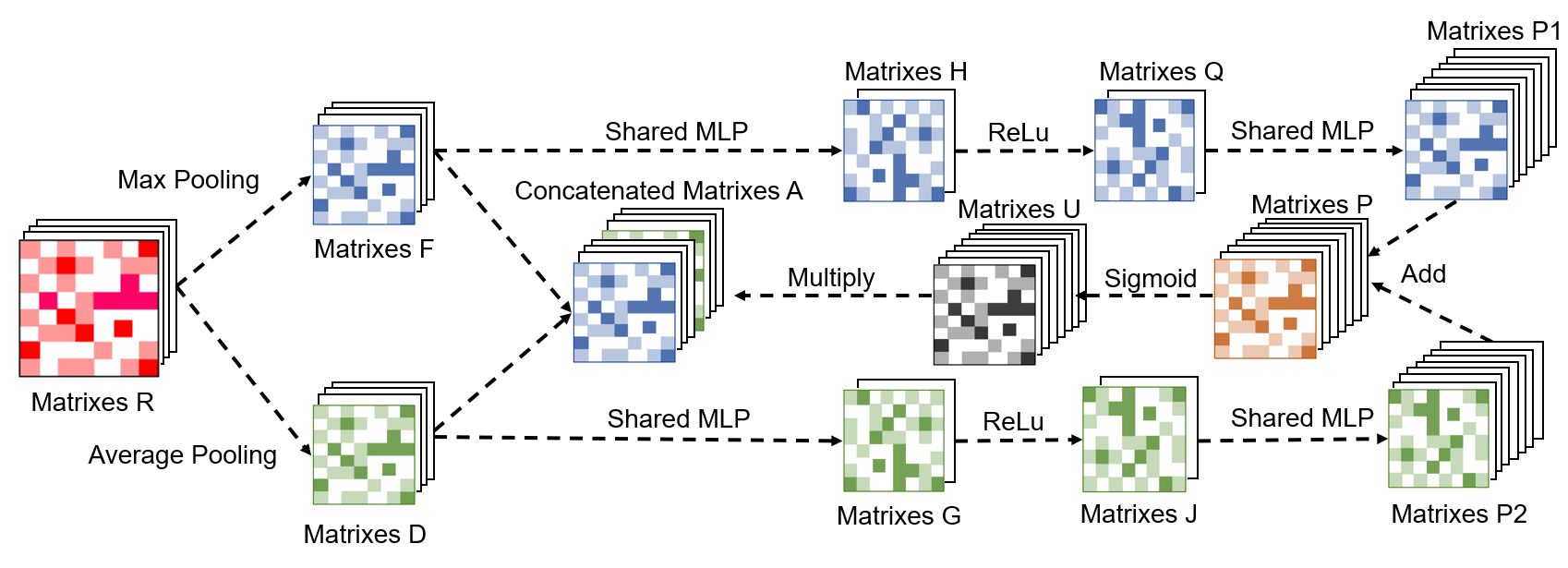

**Figure S1.** The designing of the new self-attention module in this work.

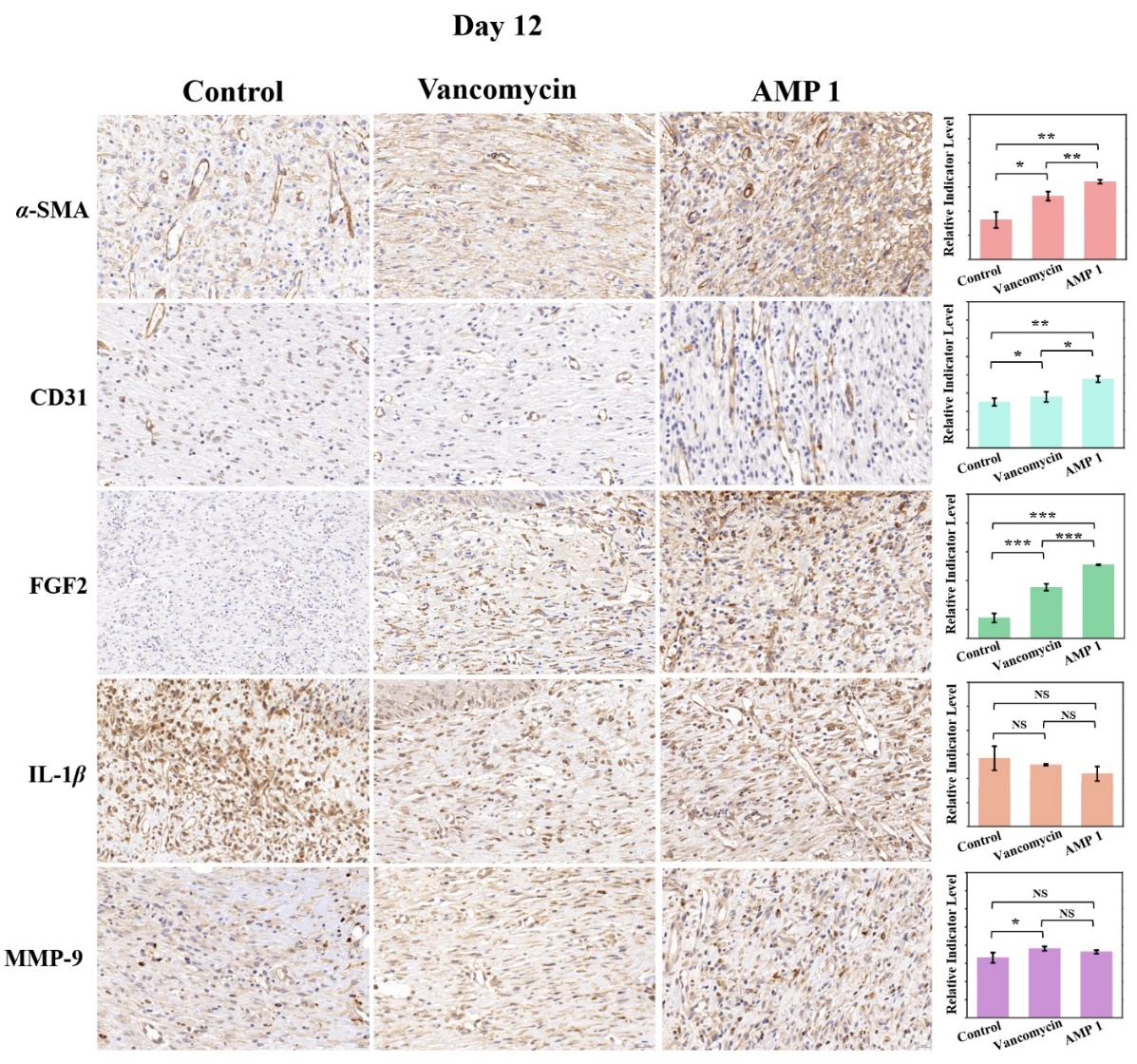

**Figure S2.** The representative histological images selected from repeated experiments for indicating *α*-SMA, CD31, FGF2, IL-1*β*, MMP-9 levels of AMP 1 treatment group and control group at Day 12. The statistical analysis of bio-markers *α*-SMA, CD31, FGF2, IL-1*β*, MMP-9 indicators levels calculated by ImageJ software among control group, Vancomycin treatment group, and AMP 1 treatment group at Day 12 with 3 sampling replicates in each group. The overall significance was assessed by t-test (*p^*^* ≤ 0.05, *p*^**^≤0.01，*p^***^* ≤ 0.001).

**Table S1. Comparison of the average performance of different model designing strategies**

| **Method** | **Sensitivity (%)** | **Specificity (%)** | **Precision (%)** | **AUPRC** |
| --- | --- | --- | --- | --- |
| A | 87.307±2.553 | 99.820±0.026 | 92.407±1.210 | 0.947±0.002 |
| B | 84.540±2.885 | 99.033±0.208 | 69.607±4.198 | 0.837±0.012 |
| C | 83.870±1.731 | 99.640±0.125 | 85.773±3.511 | 0.904±0.007 |
| D | 85.287±3.447 | 99.653±0.091 | 83.867±7.129 | 0.923±0.005 |

**Table S2. Comparison of the average performance among frequently-used algorithms in AMPs prediction**

| **Method** | **Sensitivity (%)** | **Specificity (%)** | **Precision (%)** | **AUPRC** |
| --- | --- | --- | --- | --- |
| Random Forest | 84.373±0.451 | 99.693±0.006 | 87.657±0.403 | 0.920±0.002 |
| XGBoost | 89.663±0.844 | 99.753±0.021 | 90.300±0.860 | 0.947±0.004 |
| Decision Tree | 87.170 | 98.250 | 56.260 | 0.927 |
| Linear Regression | 47.370 | 99.560 | 73.400 | 0.608 |
| Dense-Net (CNN) | 85.083±2.857 | 99.200±0.470 | 74.567±10.058 | 0.864±0.014 |
| LSTM | 82.997±4.967 | 99.277±0.331 | 75.577±8.092 | 0.869±0.025 |
| Transformer | 79.193±3.974 | 99.220±0.131 | 72.577±2.582 | 0.8175±0.005 |

Notes: For Decision Tree and Linear Regression, the training results will not be changed if the model parameters are fixed. The well-trained model files of each comparison model and their highest/average performances data generated by 3 independent trainings can be found in “Model” directory at the GitHub link.
