## Supplementary figures and images for "The discovery of antimicrobial peptides from the gut microbiome of cockroach *Blattella germanica* using deep learning pipeline"

### Figure1.tiff

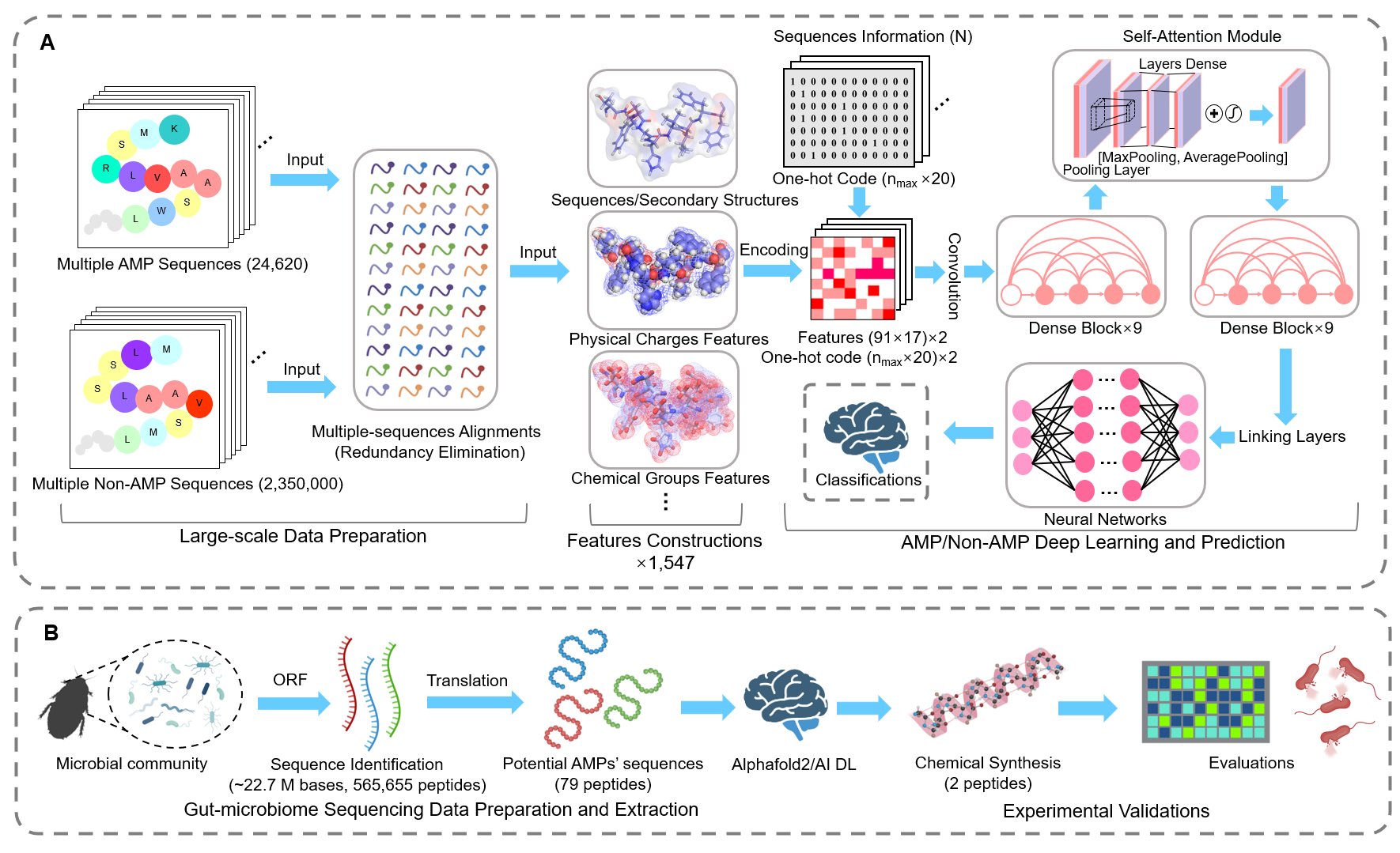

### Figure2.tiff

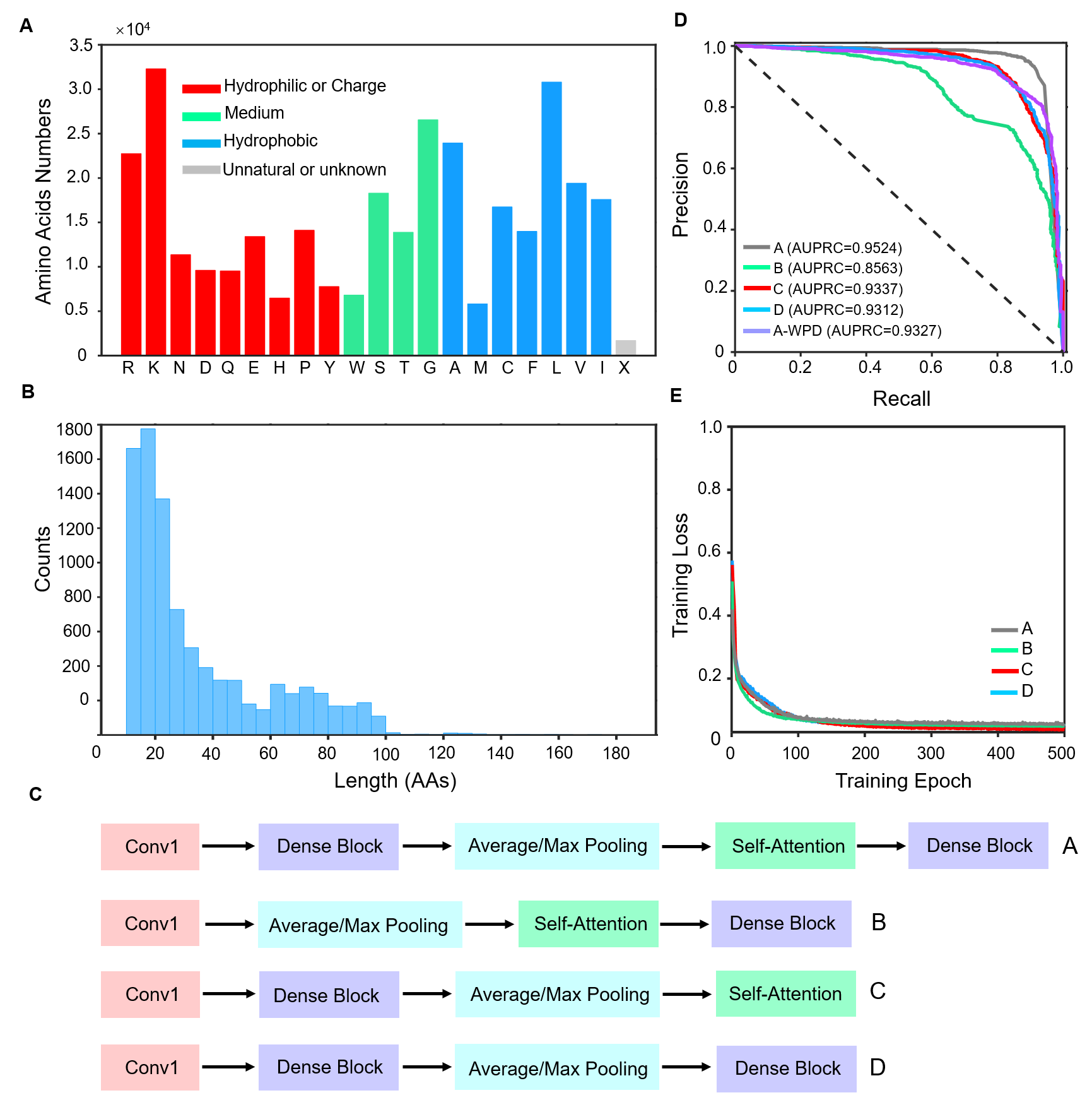

### Figure3.tiff

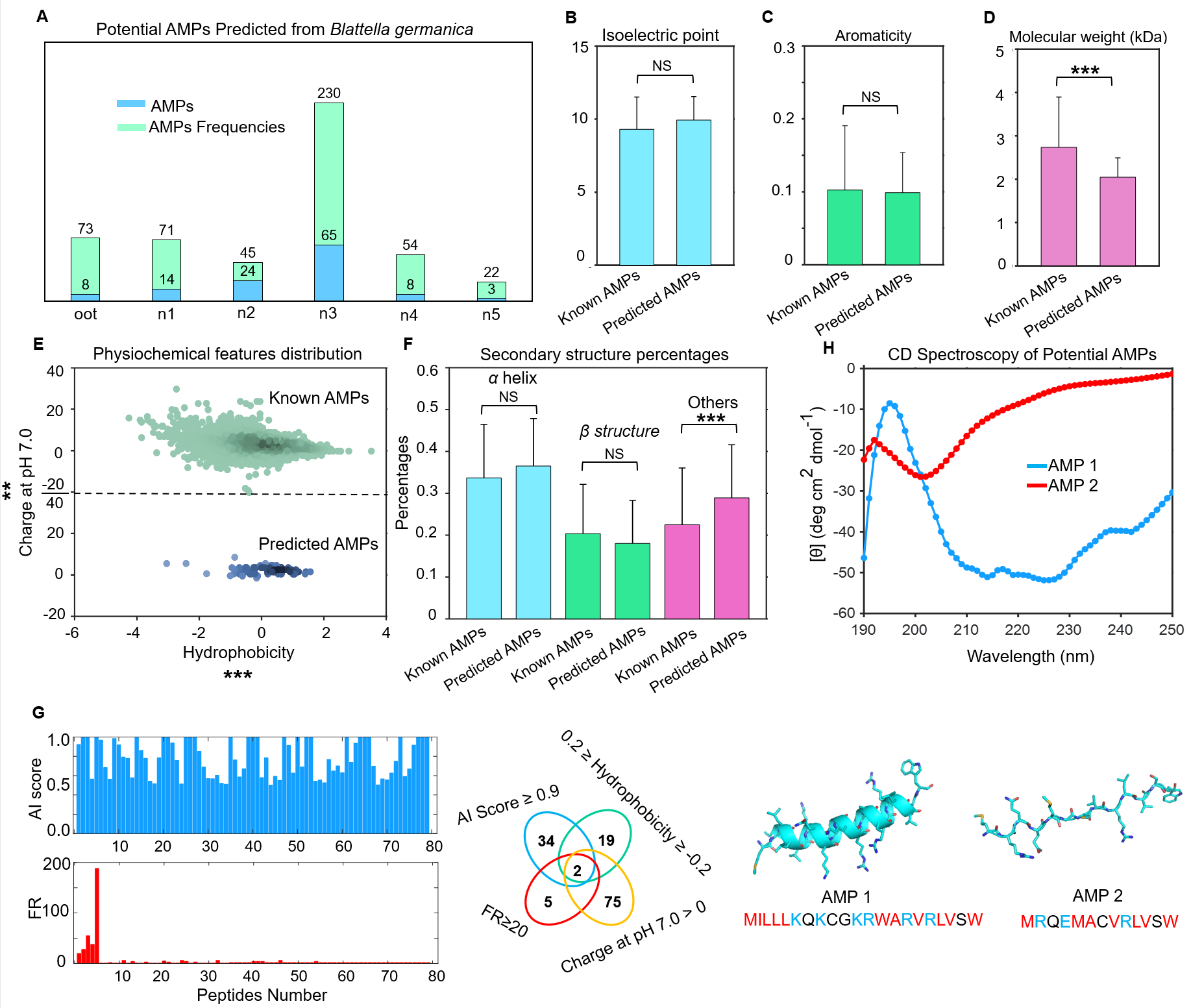

### Figure4.tiff

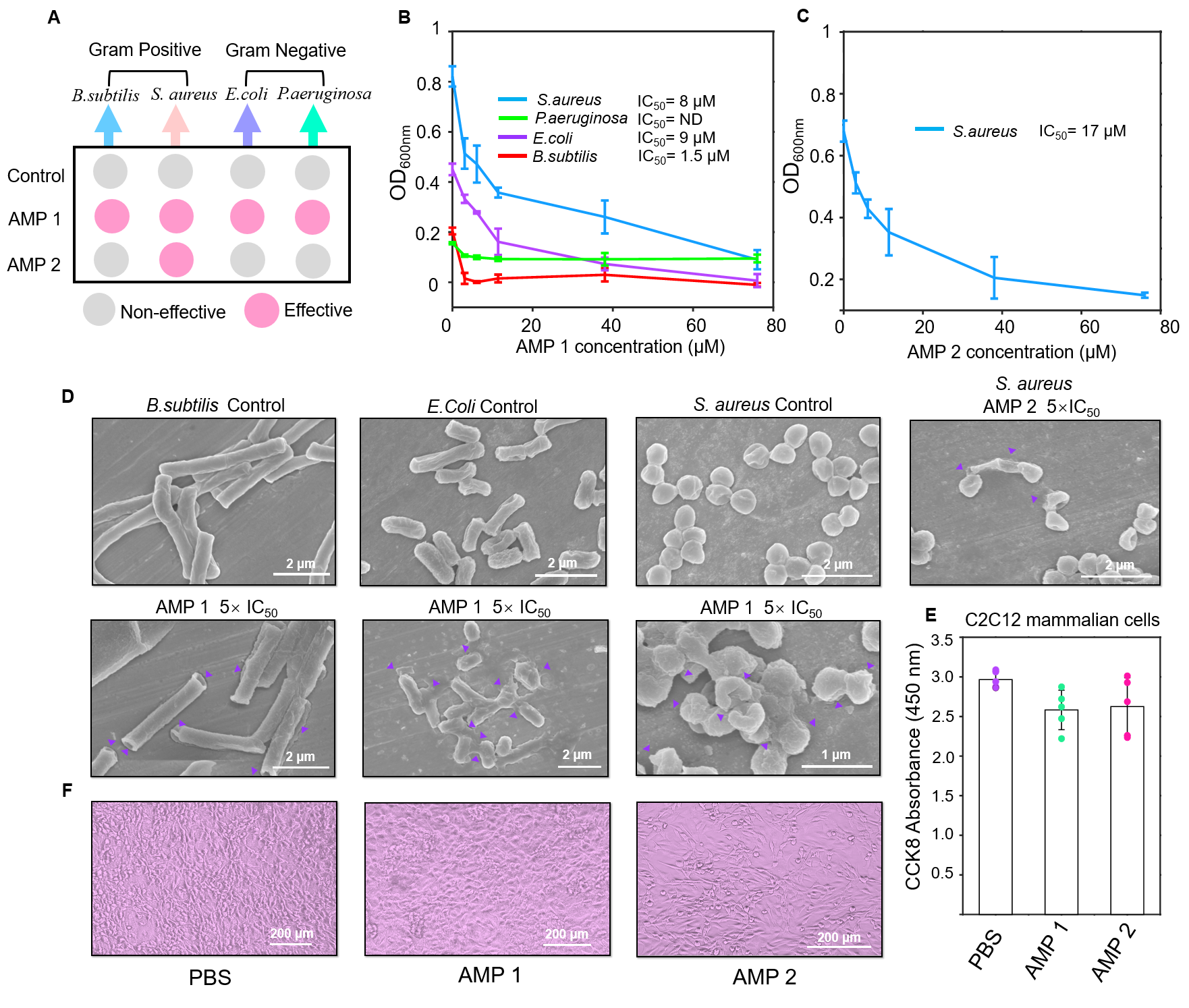

### Figure5.tiff

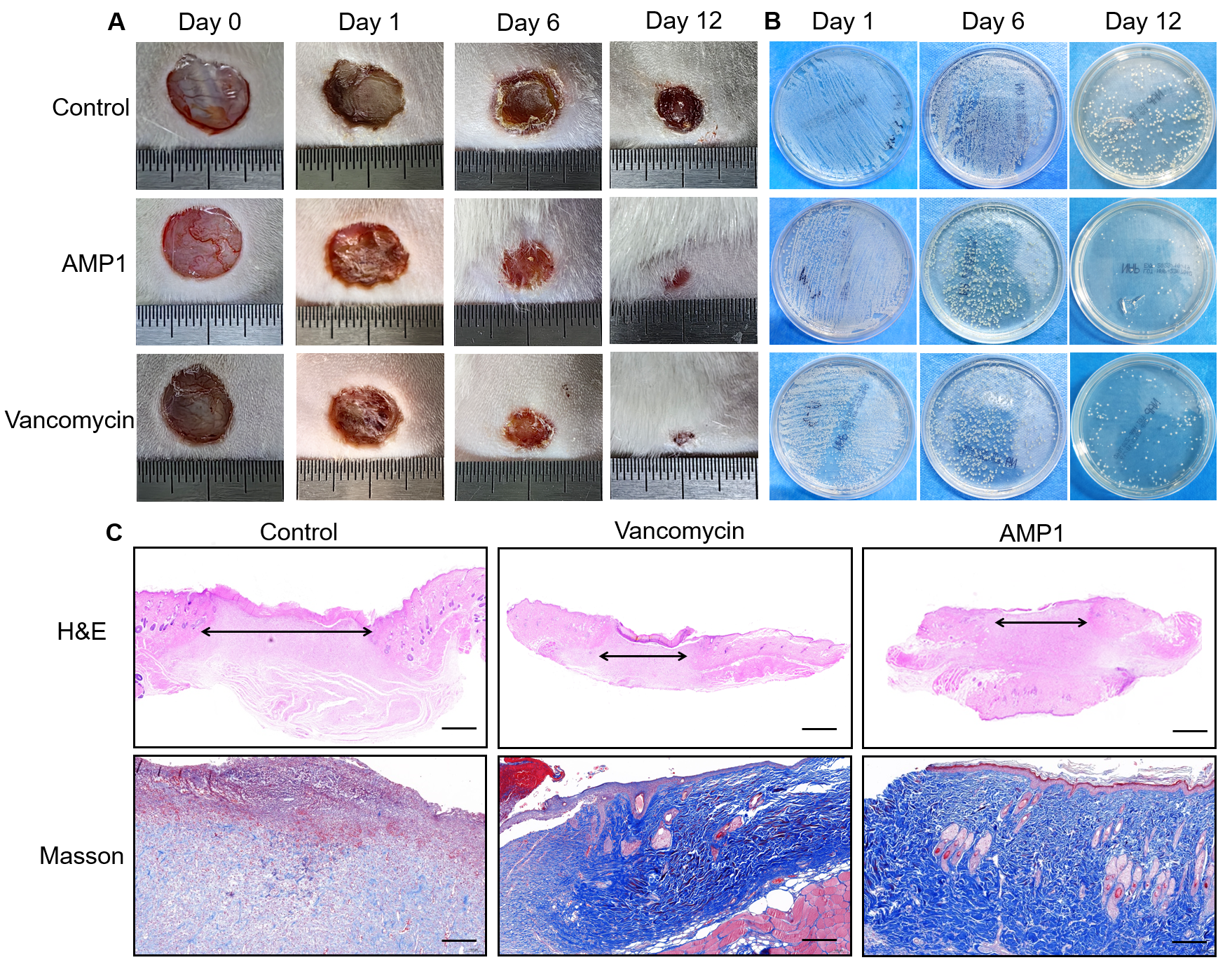

### Figure6.tiff

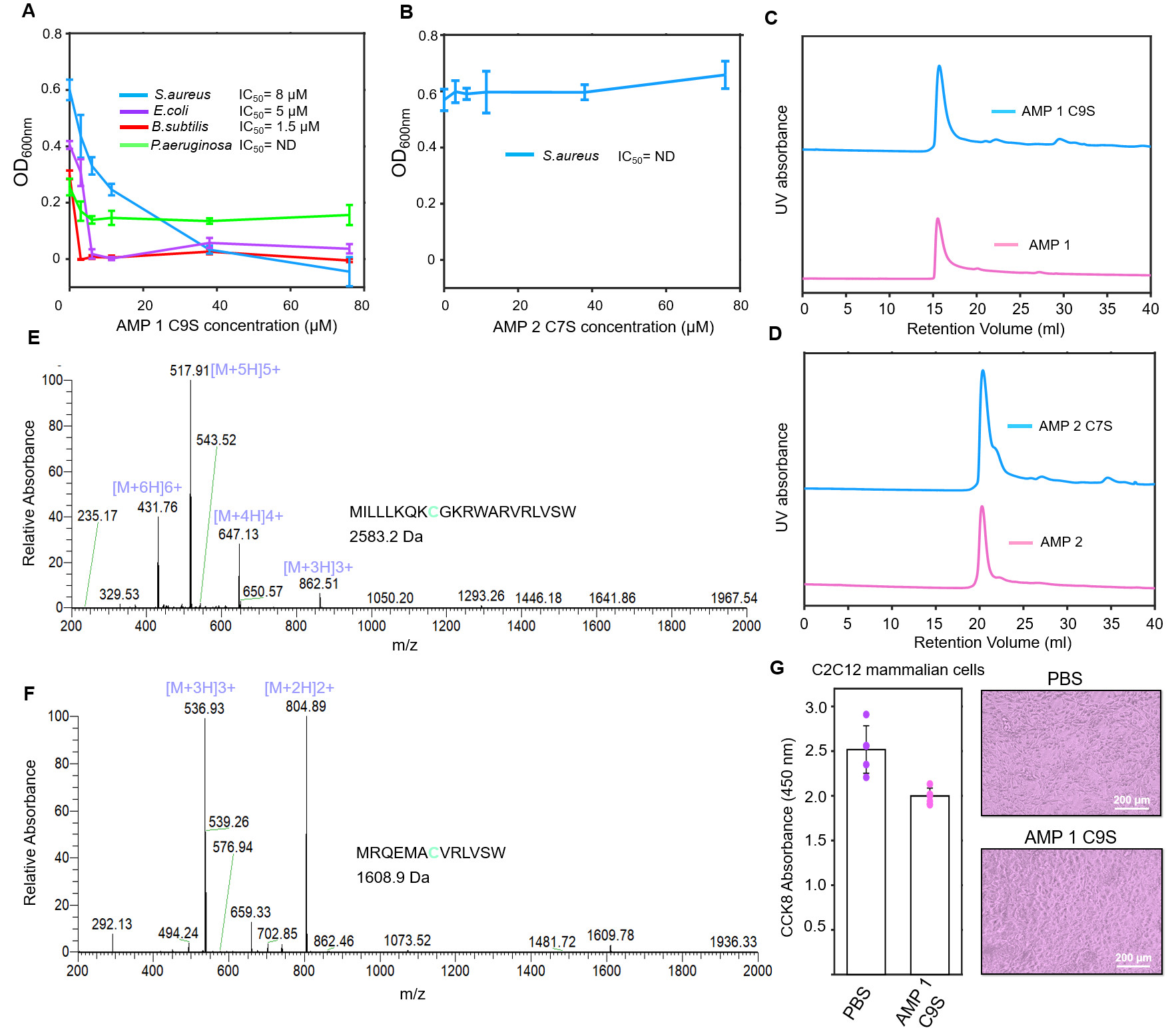

### Figure S2.tiff

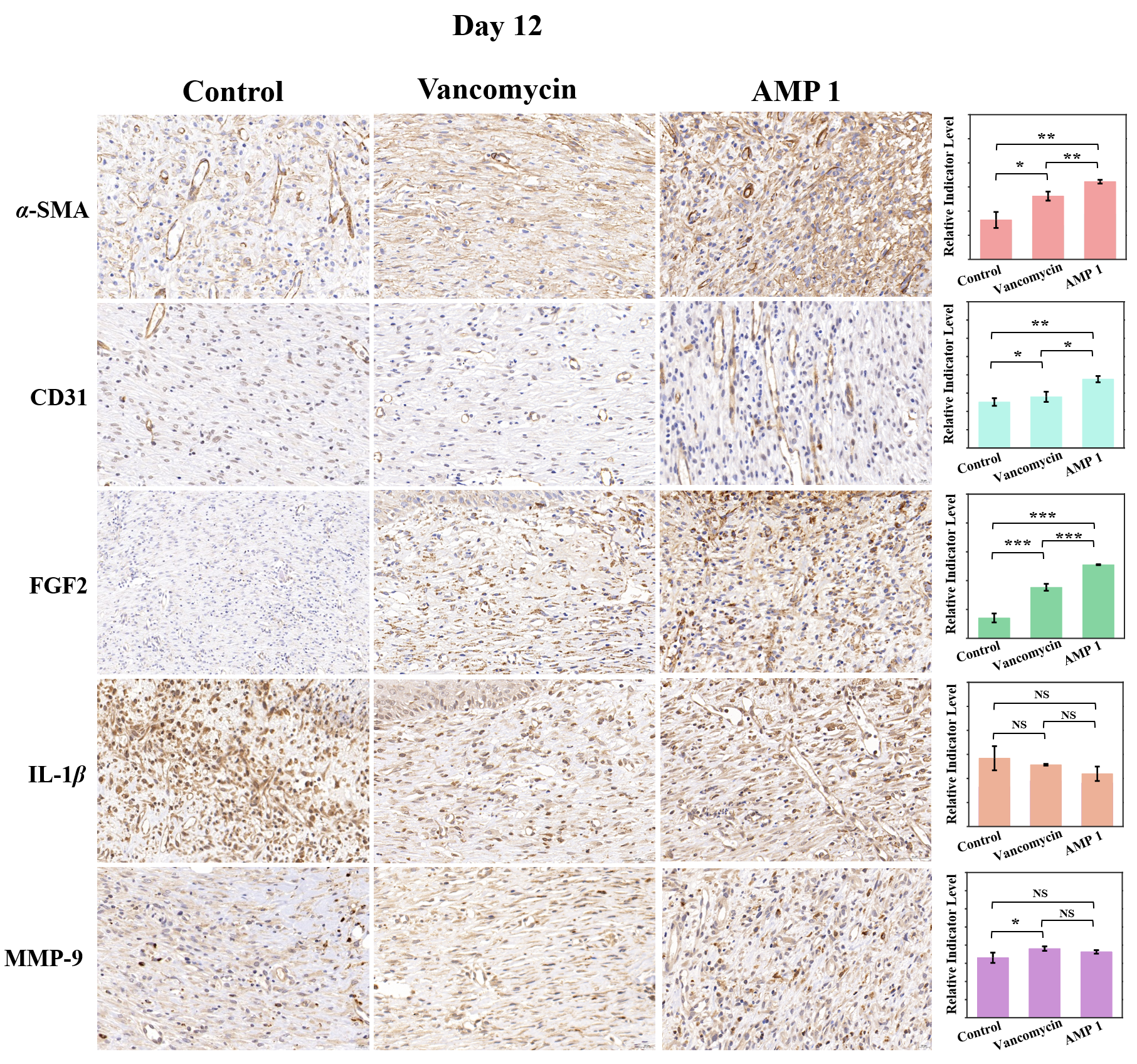

### FigureS1.tiff

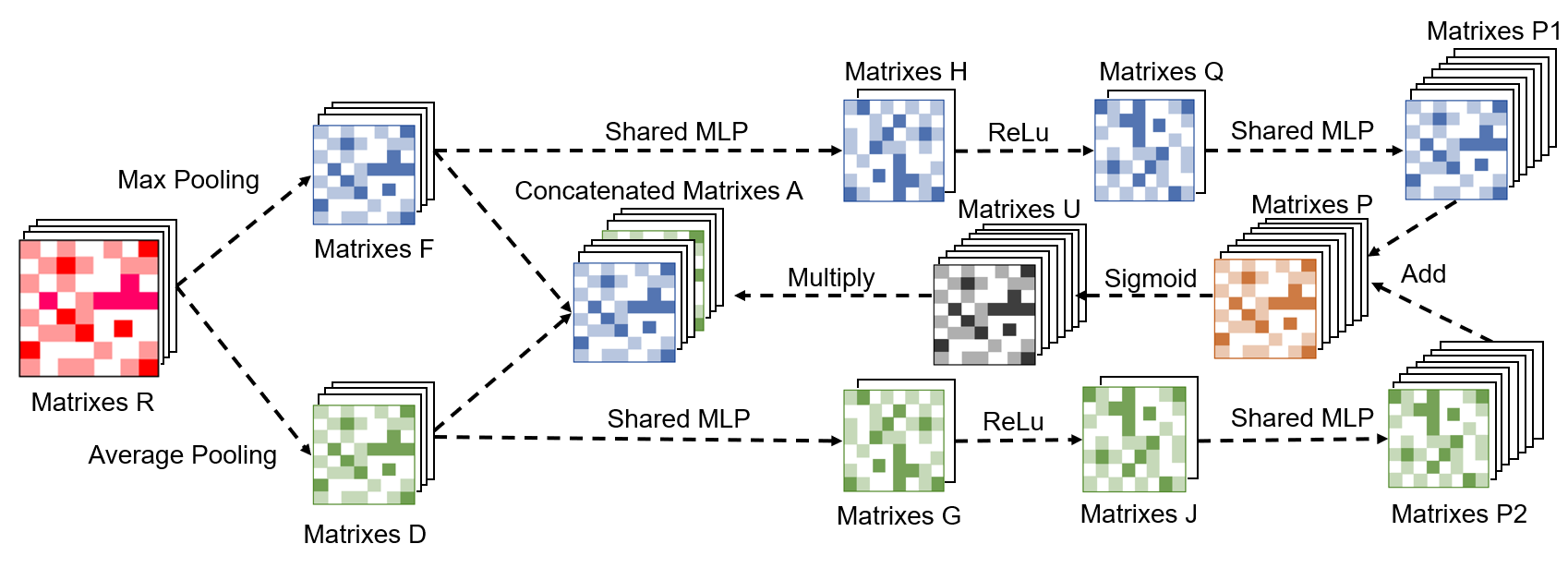

### Graphic Abstract.tiff

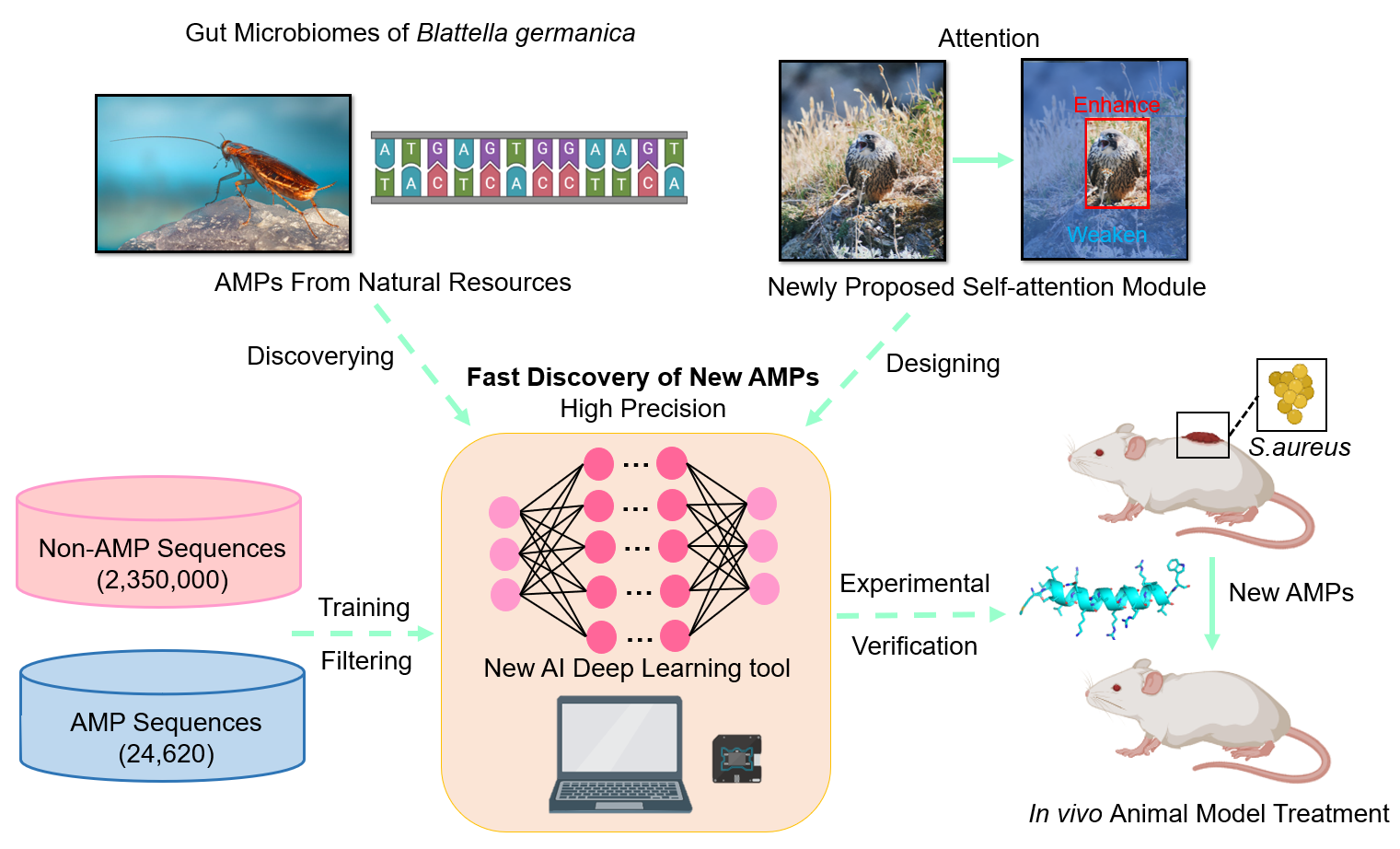
